# Dissecting Force Transmission across SUN Proteins Using Nuclear Tension Sensors

**DOI:** 10.1101/2025.02.17.638756

**Authors:** Jinfeng Wang, Juhui Qiu, Shaoying Lu, Saiyang Ding, Fangning Xu, Yanfeng Luo, Jing Zhou, Qin Peng

**Affiliations:** Institute of Systems and Physical Biology, Shenzhen Bay Laboratory, Shenzhen, 518132, P. R. China; Key Laboratory of Biorheological Science and Technology, College of Bioengineering, Chongqing University, Chongqing, 400030, China; Viterbi School of Engineering, University of Southern California, Los Angeles, CA 90089, USA; State Key Laboratory of Crop Stress Biology for Arid Areas, College of Life Sciences, Northwest A&F University, Yangling, 712100, Shaanxi, China; Department of Physiology and Pathophysiology, School of Basic Medical Sciences, State Key Laboratory of Vascular Homeostasis and Remodeling, Department of Cardiology and Institute of Vascular Medicine, Peking University Third Hospital, National Health Commission Key Laboratory of Cardiovascular Molecular Biology and Regulatory Peptides, Beijing Key Laboratory of Cardiovascular Receptors Research, Peking University, Beijing, 100191, China

## Abstract

Forces on the linker of the nucleoskeleton and cytoskeleton (LINC) complex control nuclear mechano-sensing and mechano-adaptation. However, the force transmission dynamics across the LINC complex are not fully understood, mainly because of the lack of imaging tools. We developed a set of genetically-encoded fluorescence resonance energy transfer (FRET)-based nuclear tension sensors (NuTS) that measure tension forces across SUN1/2 proteins in living cells with high sensitivity. SUN2-based NuTS (NuTS2) responded rapidly to mechanical changes in cell contractility and matrix stiffness. Notably, NuTS2 dynamically showed force transmission with high spatiotemporal resolution during cell adhesion, migration and squeeze. We also used NuTS2 to monitor tension force changes as the notochord matures in zebrafish development. NuTS2 detected a gradient tension force that increased from the posterior tail buds to the anterior as vacuoles expanded in notochord cells. Force reduction affected notochord maturation and zebrafish embryo development. Our results provide a biophysical cue for dissecting how force transmission on SUN2 protein regulates embryo development.

**Teaser:** Nuclear tension sensor (NuTS) was developed for direct force measurement across SUN proteins in living cells and zebrafish.

## Introduction

Mechanical forces generated by matrix stiffness (*1–3*), fluid shear (*4*), and mechanical stretch (*5, 6*), are critical regulators of numerous physiological and pathological processes, including embryonic development (*7–9*), tumor metastasis (*10, 11*), and cardiovascular lesions (*12, 13*). External forces can be directly transmitted by membrane-embedded receptors to the nucleus through the cytoskeleton and the linker of nucleoskeleton and cytoskeleton (LINC) complex (*14, 15*), thus modulating nuclear mechano-sensing and mechano-adaptation (*5, 16*). Force changes on nucleus enable chromatin remodeling at the nuclear periphery, thereby altering gene expression and cell fate (*5, 17–24*). Notably, strong forces on the nucleus can cause nuclear membrane rupture and chromatin damage, leading to pathological changes (*25, 26*). Therefore, determining the magnitude of forces transmitted across the LINC complex into the nucleus under physiological conditions is essential to understand the mechanical strength of the nucleus in maintaining cell homeostasis. However, measuring forces on the nucleus in living cells remains a substantial challenge.

The LINC complex, consisted of nesprin and SUN proteins, transmits mechanical forces from the cytoskeleton into the nucleus (*27*). Nesprin proteins, containing the KASH domain, span the outer nuclear membrane and link the cytoskeleton with SUN proteins, which locate on the inner nuclear membrane and connect to the nuclear lamina proteins (*28*). SUN proteins link to the nucleoskeleton lamin A/C through their N-terminal LMNA binding domain (*29*), while their C-terminal SUN domain links to the C-terminal KASH domain of nesprin embedded in the outer nuclear membrane (*30, 31*). Therefore, nesprin and SUN proteins are central to the force transmission as they form the bridge across the nuclear envelope.

Fluorescence resonance energy transfer (FRET)-based tension sensors (TS) have been developed to measure piconewton (pN)-scale forces and determine forces on the molecules responsible for mechanotransmission from cell membrane to the nuclear surface in living cells (*20, 32–36*). FRET-based TS typically integrates two fluorescent proteins connected by a mechanosensitive tension linker into the force-transmission molecules (*37*), such as talin (*33, 35*), vinculin (*32*), and nesprin (*34*). Forces stretch the tension linker to increase the distance in between the two fluorescent proteins, reducing FRET through either elongating or unfolding the linker peptide. For example, the flagelliform peptide linker F40 behaves like a molecular spring, gradually elongating when forces between 1 to 6 pN are applied (*32*). Other tension linkers, such as the ferredoxin-linker (FL) linker (*35*), villin headpiece peptide (HP) (*33*), and stabilized HP peptide (HPst) (*33*), are single-domain linkers with a digital force response. Currently, average forces transmitted from the cytoskeleton to the outer nuclear membrane protein nesprin have been measured using nesprin-TS with F40 (*20, 34*), and forces on nuclear lamina network have also been measured by lamin A tension FRET biosensor (*38*). However, there are no methods to directly detect the dynamics of forces on SUN proteins, which connect nesprin and lamin A, and are essential for measuring the forces actually transmitted into the nucleus through LINC complex.

In this study, we developed FRET-based nuclear tension sensors (NuTS) to measure forces across SUN proteins in single living cells and live zebrafish embryos. We identified that NuTS with coiled-coil structure and SUN domain at C terminus gave the most sensitive responses upon screening of different length of SUN domain and inhibitors of cell contractility. We named them as NuTS1 and NuTS2 for SUN1 and SUN2 force measurement, respectively. Both NuTS1 and NuTS2 can detect different nuclear force changes globally and locally with high spatiotemporal resolution, with NuTS2 exhibiting more force changes than NuTS1 in living cells. We applied NuTS2 to monitor force changes during notochord maturation in the zebrafish embryos, and observed a gradient force increases from posterior tail buds to the anterior as vacuoles enlarged in notochord cells. Our results provide direct evidence of the correlation between nuclear forces and notochord maturation in regulating embryo development.

## Results

### Generating and screening of NuTS1 and NuTS2

It is well known that among the five different SUN domain containing proteins, SUN1 and SUN2 are ubiquitously expressed and partially redundant (*39*). To measure the mechanical force transmission across the inner nuclear membranes through SUN1/2 protein in living cells, following the existing FRET-force module design principle (*32, 34, 40*), we constructed SUN1/2 NuTS biosensors with five parts in general: SUN’s N-terminal LMNA-binding domain and transmembrane peptides (TM), an enhanced cyan FP (ECFP) and a yellow FP variant (YPet) for FRET, F40 tension linker (1-6 pN) between ECFP and YPet, and coiled-coil structure (CC) and SUN domain from SUN proteins as C-terminus. The most important domain inside SUN domain is KASH lid, which interacts with KASH domain of nesprin (*41*). To screen if there were any other domains besides KASH lid that are essential for force transmission on SUN protein, we had three designs for NuTS, one was with KASH lid only (KL-NuTS), the other was with intact SUN domain (SD-NuTS), and another was with both coiled-coil structure and SUN domain (CSD-NuTS). As a control, Tail-Less (TL), followed the same pattern as NuTS but without any domains after ECFP, bears zero force exhibiting the highest FRET ratio. We named biosensors as NuTS1 and TL1 for SUN1, and NuTS2 and TL2 for SUN2, respectively (Fig. 1A and fig. S1). When NuTS subjects force from the upstream nesprin and downstream lamin A, the tension linker in NuTS gets stretched and the two fluorescent proteins apart from each other, resulting in the reduction in FRET signal. Therefore, FRET ratio is negatively correlated with mechanical force, and the lower FRET ratio corresponds to the higher tension force on SUN proteins (Fig. 1B). With their overexpression in wild-type (WT) or *SUN1/2* knockout (KO) cells, both NuTS1 and NuTS2 constructs properly localized on the nuclear membrane (NM) with lower FRET ratios, compared to TL1 or TL2 (Fig. 1, C and F, and fig. S2). In both WT and SUN KO cells, KL-NuTS1 exhibited the lowest FRET basal level and the most FRET change on SUN1 protein, while KL-NuTS2 and CSD-NuTS2 behaved the same for force measurement on SUN 2 protein, compared to TL groups (Fig. 1, D, E, G and H), in both HeLa and C2C12 cell lines. Since there was no difference in FRET ratio between WT and knockout cells (Fig. 1, C and F), we used WT cells for biosensor characterization and application in the following experiments. Overall, the localization and function of NuTS is independent from cell type and endogenous SUN protein level, but further characterization is required to find the best design for both NuTS1 and NuTS2 in response to nuclear force changes.

**Fig. 1.**
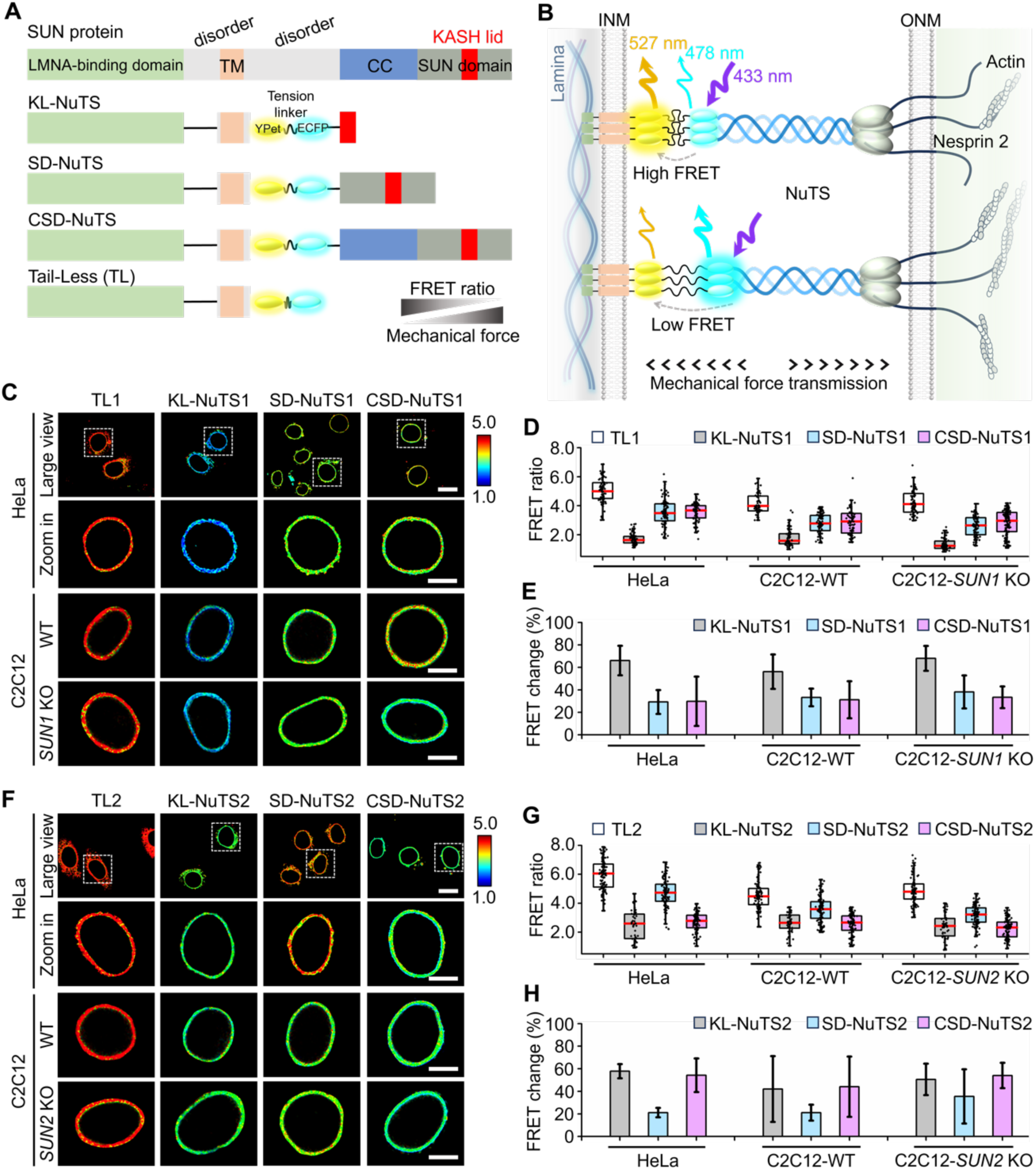
Designation and domain screening of FRET-based nuclear tension sensors (NuTS). (**A**) Scheme of SUN proteins, three designs of the SUN-based nuclear tension sensor (NuTS), and truncated tail-less (TL). (**B**) The working mechanism of the mechanical sensing of NuTS. FRET ratio is negatively correlated with mechanical force. Low FRET of NuTS represents large force transmission on SUN protein, vice versa. (**C**) Representative FRET ratiometric images of TL1 and three designs of NuTS1 transiently expressed in HeLa-WT, C2C12-WT, and C2C12-*SUN1* KO cells, respectively. Scale bars, 20 μm for large view, and 10 μm for zoom in. (**D**) The quantified FRET ratio (Mean ± SEM) of TL1 and series NuTS1 from (**C**). n = 81, 76, 101, 81 (HeLa), n = 59, 62, 93, 85 (C2C12-WT), and n = 82, 65, 99, 119 (C2C12-*SUN1* KO). (**E**) Fold changes of three designs of NuTS1 calculated by (TL1-NuTS1)/TL1*100% from (**D**). KL-NuTS1 showed the most FRET ratio change compared to TL1 in three types of cells. (**F**) Representative FRET ratiometric images of TL2 and three designs of NuTS2 transiently expressed in HeLa-WT, C2C12-WT, and C2C12-*SUN2* KO cells, respectively. Scale bars, 20 μm for large view, and 10 μm for zoom in. (**G**) The quantified FRET ratio (Mean ± SEM) of TL2 and series NuTS2 from (**F**). n = 114, 48, 116, 104 (HeLa), n = 103, 54, 105, 104 (C2C12-WT), and n = 88, 54, 102, 101 (C2C12-*SUN2* KO). (**H**) Fold changes of three designs of NuTS2 calculated by (TL2-NuTS2)/TL2*100% from (**G**). KL-NuTS2 and CSD-NuTS2 displayed higher FRET ratio changes than SD-NuTS2 in all type of cells. Three biological replicates were performed for all experiments, respectively. Ordinary one-way ANOVA Tukey’s multiple comparisons.

### NuTS1 and NuTS2 respond to force changes by cytoskeleton-mediated cell contractility

To further evaluate the sensitivity of NuTS, we treated cells transiently expressing NuTS with small molecules to change cell contractility. Myosin inhibitor Blebbistatin (Blebb) (*42, 43*) and actin polymerization inhibitor Cytochalasin D (Cyto D) (*44, 45*) were used to decrease the cell contractility. Significant nuclear membrane wrinkling was observed upon Blebb and Cyto D treatment (fig. S3), indicating the reduction of cell contractility by small molecules effectively. Compared to DMSO group, CSD-NuTS design showed the most FRET changes in response to Blebb and Cyto D for both NuTS1 and NuTS2 (Fig. 2, A-F). The FRET changes of CSD-NuTS2 reached approximately 60% while CSD-NuTS1 reached 30% maximumly, implying SUN2 plays more dominant role than SUN1 in transducing forces from cytoskeleton to nucleus. It was worth noting that, KL-NuTS design had the shortest length among three designs, so likely, its F40 linker got extended the most when KL-NuTS spanned the 30-50 nm of the nuclear envelop lumen, resulting in constitutive high tension on the sensor with lowest FRET always (Fig. 2, A-F), no matter there was contraction force from cytoskeleton or not. In addition, calyculin A (Caly A), a potent inhibitor of myosin light chain phosphatase to enhance F-actin sliding by over-activating myosin (*46*), was used to increase cell contractility. The results showed that FRET ratio from all the designs of NuTS1 and NuTS2 did not show any notable change (Fig. 2, A-F). We speculated that the force on SUN proteins under calyculin A treatment might have exceeded the upper limit of F40 tension linker, so none of the NuTS designs whichever measured by F40 linker responded. Taken together, the most optimized design for both SUN1 and SUN2 force measurement was the same, NuTS with coiled-coil structure and SUN domain at C terminus (CSD-NuTS), which were defined as NuTS1 and NuTS2 in this study. Moreover, time-lapse imaging demonstrated the FRET dynamics of NuTS2 in monitoring force changes on SUN2 protein upon inhibiting actin polymerization by Cyto D or actomyosin phosphorylation by Y-27632 (Fig. 2, G-J, movie S1 and S2). The FRET increased rapidly within 15 min upon Cyto D or Y-27632 addition while no changes in TL2 groups (Fig. 2, H and J), although the FRET increase was not as much as the FRET changes in Fig. 2F likely due to the different ways for sensor expression. These data demonstrated that NuTS1 and NuTS2 can effectively and sensitively respond to the actin-mediated force changes.

**Fig. 2.**
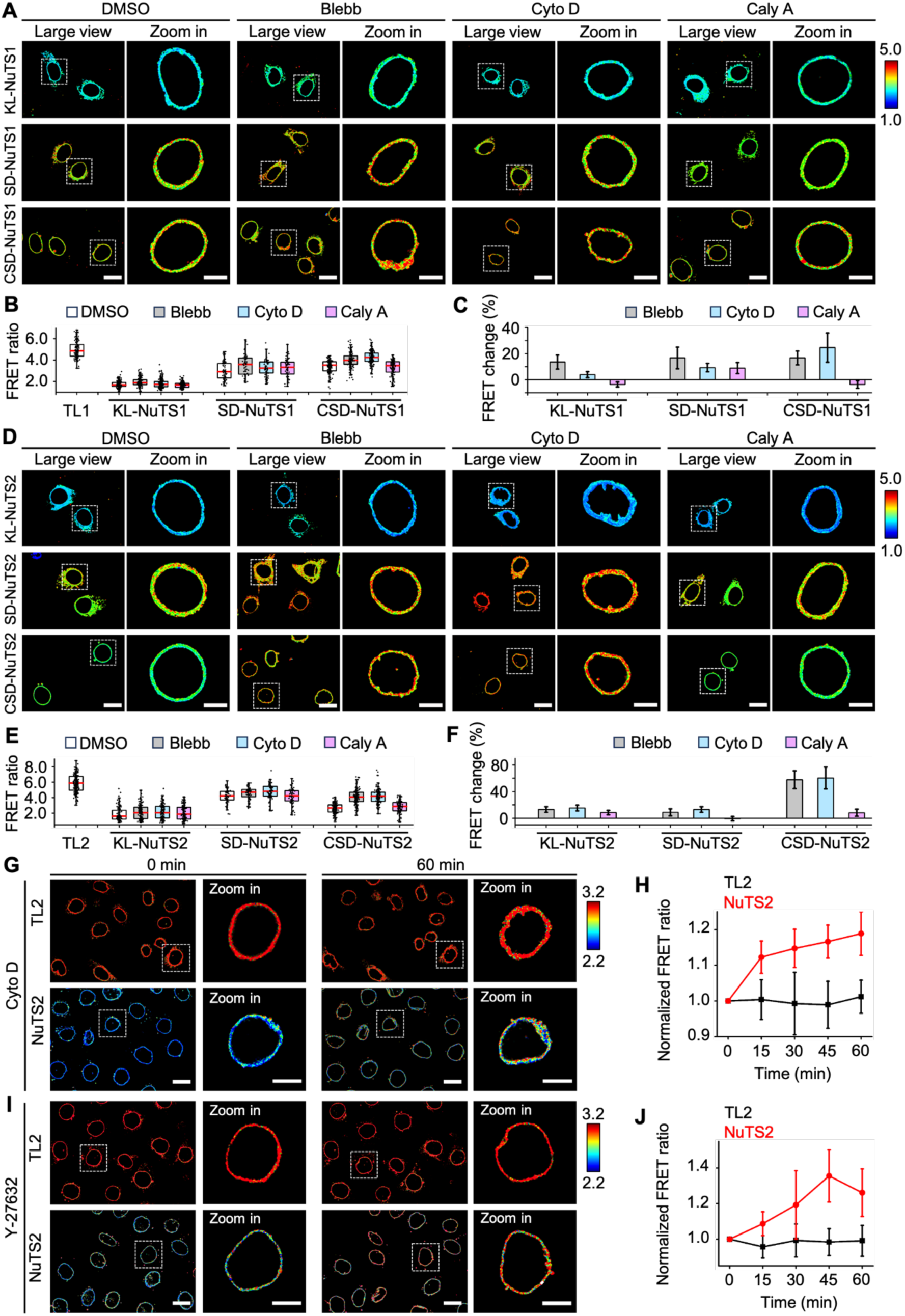
Characterization the specificity of NuTS in response to force changes by the contraction of cytoskeleton. (**A**) Representative FRET ratiometric images of three types of NuTS1 transiently expressed in HeLa cells. Cells were treated with 1‰ DMSO (control), 30 μM Blebb, 1 μM Cyto D, and 1 nM Caly A, respectively. Scale bars, 20 μm for large view, and 10 μm for zoom in. (**B**) The quantified FRET ratio (Mean ± SEM) of TL1 and series NuTS1 from (**A**). n = 217 (TL1), n = 184, 162, 202, 188 (KL-NuTS1), n = 72, 72, 72, 72 (SD-NuTS1), and n = 154, 150, 152, 150 (CSD-NuTS1). (**C**) Fold changes of series NuTS1 calculated by (Drug-DMSO)/DMSO*100% from (**B**). (**D**) Representative FRET ratiometric images of three types of NuTS2 transiently expressed in HeLa cells. Cells were treated with the same conditions as NuTS1 in (**A**). Scale bars, 20 μm for large view, and 10 μm for zoom in. (**E**) The quantified FRET ratio (Mean ± SEM) of TL2 and series NuTS2 from (**D**). n = 234 (TL2), n = 121, 127, 127, 133 (KL-NuTS2), n = 74, 73, 74, 73 (SD-NuTS2), and n = 162, 140, 149, 140 (CSD-NuTS2). (**F**) Fold changes of series NuTS2 calculated by (DrugDMSO)/DMSO*100% from (**E**). (**G**)(**I**) Time-lapse images showed the dynamic FRET changes of TL2 and NuTS2 in HeLa that were treated with 1 μM Cyto D or 10 μM Y-27632, respectively. Time interval was 15 min/frame. Scale bars, 20 μm for large view, and 10 μm for Zoom in. (**H**)(**J**) The normalized FRET ratio (Mean ± SEM) of TL2 and NuTS2 from (**G**)(**I**). n= 15, 7 (**H**), and n = 12, 15 (**J**). Three biological replicates were performed for all experiments, respectively. Ordinary one-way ANOVA Tukey’s multiple comparisons.

### Characterization the specificity of force transmission on NuTS1 and NuTS2

As an integral part of the LINC complex, SUN proteins transmit mechanical forces from cytoskeleton to nuclear lamina across the nuclear envelope (*47*). Next, we further characterized the specificity of the force transmission on NuTS1 and NuTS2. Nesprin is the upstream connection of SUN proteins to the cytoskeleton, straightforwardly, we tested NuTS1 and NuTS2 in the *SYNE2* knockout C2C12 cells. Our results showed that NuTS2 had significant increase of FRET signal in *SYNE2* KO cells compared to WT cells (Fig. 3, A-C, and fig. S4A), while NuTS1 had no significant changes in response to *SYNE2* KO (fig. S4, B and C). Lamin A is the other anchor of SUN protein, so we wondered how NuTS1 and NuTS2 responded to the loss of lamin A at downstream. There was much higher FRET ratio observed with NuTS2 in *LMNA* KO C2C12 than WT cells (Fig. 3, D-F, and fig. S5A), demonstrating its specificity and sensitivity to the connection of lamina network. However, NuTS1 didn’t respond much in either *LMNA* KO or *LMNB1* KO cells, compared to WT cells (fig. S5, A-C), indicating tension on the SUN1 sensor is not sensitive to the loss of lamin A and lamin B1. It was reported that the specialized SUN: nesprin pair is required for their function (*48*), implying SUN1 primarily interacts with nesprin 1 but not nesprin 2 in force transmission function, a possible reason that NuTS1 did not respond much in *SYNE2* KO cells. As for the downstream connection of SUN1, it interacts with a few nuclear membrane proteins, including lamin A (*29*), Nup153 (*49, 50*), and emerin (*51*), to not only transmit forces but also play other roles in stabilizing nuclear pore complex and association with other inner nuclear protein. Therefore, an alternative explanation for NuTS1 not sensitive to *LMNA* depletion is that, *LMNA* depletion may not sufficiently affect the state of SUN1.

**Fig. 3.**
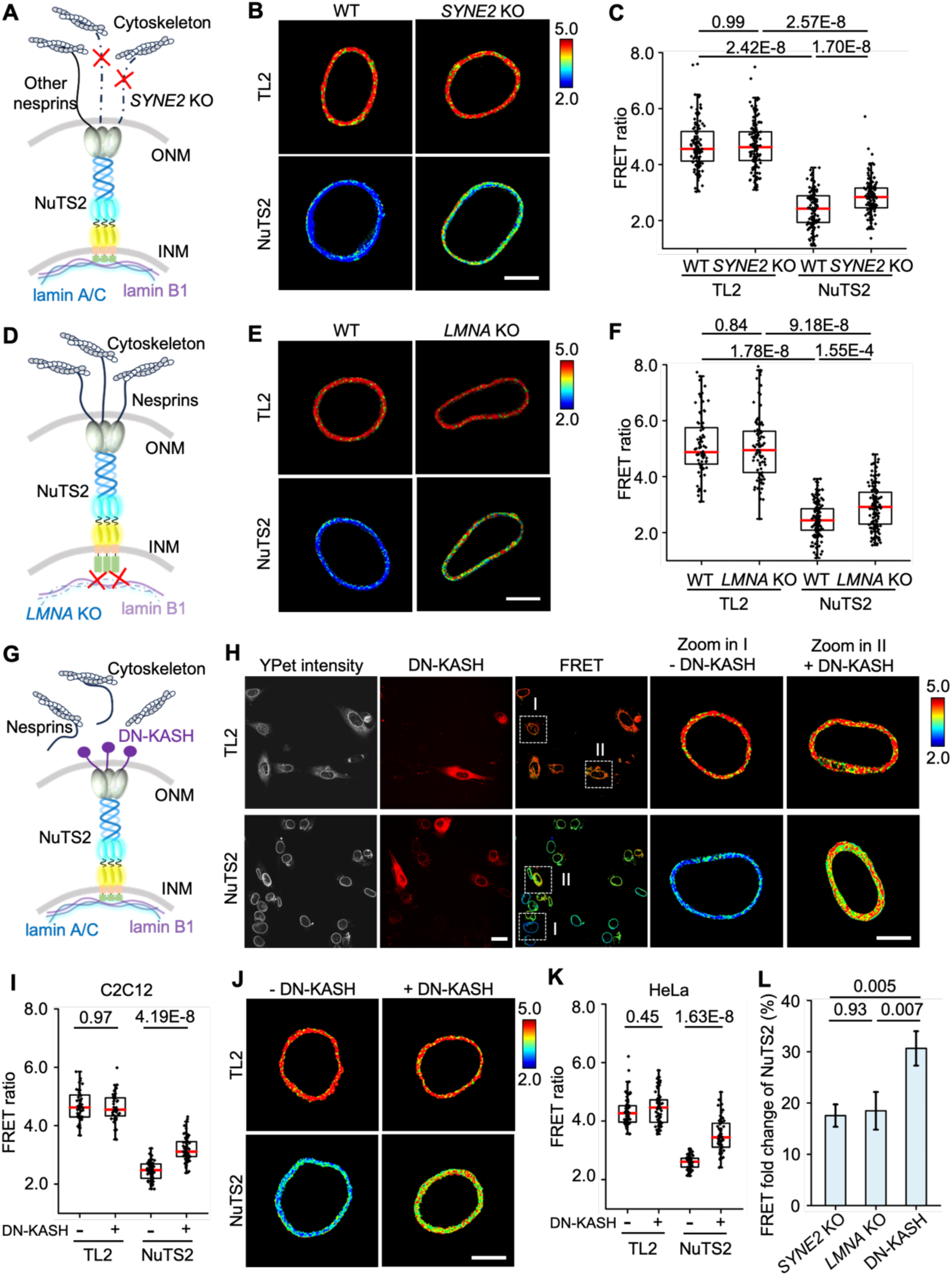
NuTS detects force transmission through intact LINC complex from cytoskeleton to nucleus. (**A**) Scheme of mechanical force transmission in *SYNE2* KO C2C12. (**B**) Representative FRET ratiometric images of TL2 and NuTS2 transiently expressed in *SYNE2* KO and WT C2C12, respectively. Scale bars, 10 μm. (**C**) The quantified FRET ratio (Mean ± SEM) of TL2 and NuTS2 from (**B**). n = 119, 131, 136, 133. (**D**) Scheme of mechanical force transmission in *LMNA* KO C2C12. (**E**) Representative FRET ratiometric images of TL2 and NuTS2 transiently expressed in *LMNA* KO and WT C2C12, respectively. Scale bars, 10 μm. (**F**) The quantified FRET ratio (Mean transmission on LINC complex with dominant negative KASH (DN-KASH) expressed. (**H**) Representative FRET ratiometric images from TL2 and NuTS2 stable C2C12 cells transiently expressing DN-KASH for 36 h, respectively. Scale bars, 20 μm for large view, and 10 μm for zoom in. (**I**) The quantified FRET ratio (Mean ± SEM) of TL2 and NuTS2 from (**H**). n = 51, 41, 82, 75. (**J**) Representative FRET ratiometric images from TL2 and NuTS2 stable HeLa cells transiently expressing DN-KASH, respectively. Scale bars, 10 μm. (**K**) The quantified FRET ratio (Mean ± SEM) of TL2 and NuTS2 from (**J**). n = 70, 71, 78, 73. (**L**) The fold changes of NuTS2 calculated by (*SYNE2* KO - WT)/WT*100% from (**C**), (*LMNA* KO - WT)/WT*100% from (**F**), and (DN-KASH ^(+)^ - DN-KASH (^-^))/DN-KASH (^-^) *100% from (**I**). Three biological replicates were performed for all experiments, respectively. Ordinary one-way ANOVA Tukey’s multiple comparisons.

Furthermore, disruption of the LINC complex using a dominant negative nesprin construct (DN-KASH) (*52*) substantially increased FRET ratio in WT C2C12 and HeLa cells monitored by NuTS2 (Fig. 3, G-K, and fig. S6), further indicating the specificity of NuTS2 in nucleus. We also quantitatively compared the effects of *SYNE2* depletion, *LMNA* depletion, and expression of the DN-KASH, and found out that, DN-KASH caused much more FRET increase than *SYNE2* depletion and *LMNA* depletion (Fig. 3L), indicating Nesprin-2 is not the only relevant KASH protein for force transmission to SUN proteins but other nesprins also play roles. Overall, NuTS2 behaved more sensitively to the force changes on SUN2 protein, which might imply SUN2 is the primary SUN protein sensitive to actin-mediated tension.

### NuTS responds to global force changes upon external and internal stimuli

Cells on stiff substrate can form a strong cytoskeleton to enhance cell contractility and transmit greater force to nucleus, on the contrary, less force is transmitted to nucleus on soft substrate (*53, 54*). We applied NuTS1 and NuTS2 to visualize the corresponding nuclear force transmission on PAA hydrogels with different matrix stiffness (fig. S7). Significant decreasing of FRET signals was observed on 40 kPa PAA gel compare to 0.2 kPa gel with both NuTS1 and NuTS2, and FRET got recovered upon Cyto D addition on stiff gels, while no changes on soft gels (Fig. 4, A-C, and fig. S8), suggesting that tension forces on SUN proteins increased on stiff substrate mediated by actin externally. Chromatin condensation status has been shown to affect nuclear mechanics (*55*). People have measured the mechanical properties of nuclei by using atomic force microscopy and micropipette aspiration (*55–57*). After trichostatin A (TSA) treatment, a histone deacetylase inhibitor, the stiffness of nuclei got reduced, resulting in decreasing in nuclear tension forces. To measure this, we treated cells with TSA to decondense chromatin.

**Fig. 4.**
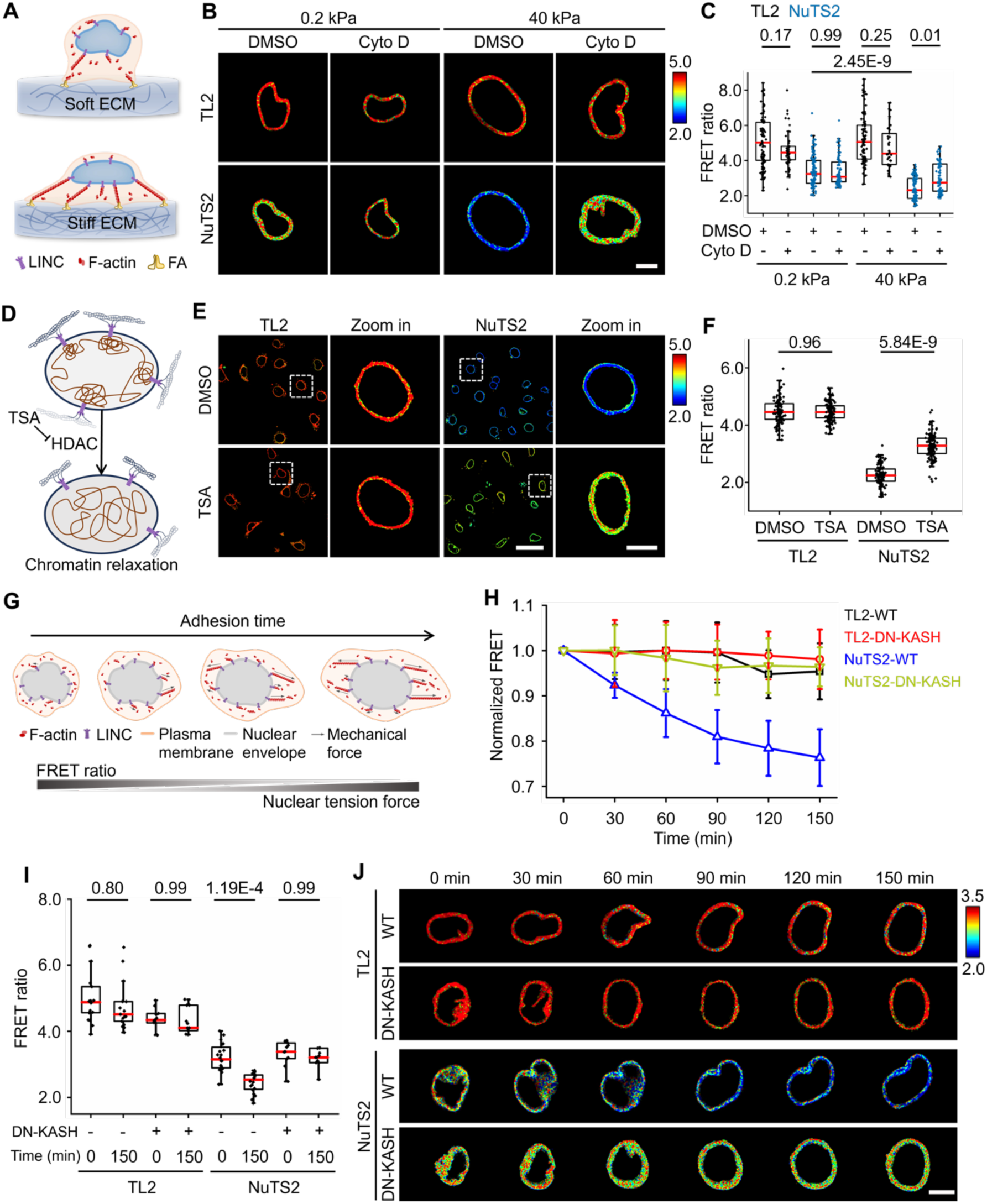
NuTS responds to global force changes from external and internal stimuli. (**A**) Scheme of ECM stiffness effect on cell contractility and nuclear mechanics. FA, focal adhesion. Cells spread more on stiff extracellular matrix (ECM). (**B**) Representative FRET ratiometric images of TL2 and NuTS2 expressed in C2C12-WT that seeded on fibronectin coated soft (0.2 kPa) and stiff (40 kPa) PAA gels, respectively. Scale bars, 10 μm. (**C**) The quantified FRET ratio (Mean ± SEM) of TL2 and NuTS2 from (**B**) n = 106, 51, 126, 80, 101, 47, 93, 84. (**D**) Schematic diagram of chromatin organization effect on nuclear tension by treated with TSA (histone deacetylase inhibitor). HDAC, histone deacetylase. (**E**) Representative FRET ratiometric images of stable HeLa cells expressing TL2 and NuTS2 with 500 nM TSA for 20 h, respectively. Scale bars, 50 μm for large view and 10 μm for zoom in. (**F**) The quantified FRET ratio (Mean ± SEM) of TL2 NuTS2 from (**E**). n = 126, 128, 119, 132. (**G**) Schematic diagram of cell adhesion process. The mechanical force transmission across nuclear membranes increases along cell spreading. (**H**) The normalized FRET ratio (Mean ± SEM) of TL2 and NuTS2 with or without DN-KASH during cell spreading. n = 17, 11, 20, 11. (**I**) The quantified FRET ratio (Mean ± SEM) of TL2 and NuTS2 before (0 min) and after (150 min) cell adhesion. n = 17, 11, 20, 11. (**J**) FRET Time-lapse images showed the FRET dynamics of TL2 and NuTS2 with or without DN-KASH in representative WT C2C12 upon seeding on glass bottom dishes. Time interval was 30 min. Scale bars, 10 μm. Three biological replicates were performed for all experiments, respectively. Ordinary one-way ANOVA Tukey’s multiple comparisons.

Compare to the control group with DMSO, our imaging results showed the significant increase of FRET ratio in HeLa stable cells expressing NuTS2 with TSA treatment (Fig. 4, D-F), implying chromatin decondensation reduces tension forces on SUN2 protein, likely due to the dissociation of relaxed chromatin from inner nuclear membrane, lose the connection to the LINC complex and become less able to resist deformation. Moreover, we noticed that individual cells displayed different levels of FRET signals (Fig. 4E). We hypothesized that cell adhesion and spreading status might cause different cytoskeleton-nucleus force transmission dynamically. We subsequently tested if NuTS reacted to the global changes in force on nucleus. We monitored cell adhesion process by using NuTS2, and observed continuous FRET ratio decreasing during cell spreading, but no significant FRET changes on TL2 (Fig. 4, G-J), indicating the increase of nuclear force transmission across SUN2 in cell adhesion. Compared to WT cells, there was no significant changes of FRET signal during cell adhesion process with removing nesprin-2 or blocking KASH binding by DN-KASH, demonstrating the FRET signal change on NuTS in WT cells is due to tension forces on the LINC complex (Fig. 4H-J, fig. S9). Meanwhile, NuTS2 detected the global reduction of nuclear forces across SUN2 protein under TGF-β1 induced epithelial to mesenchymal transition (EMT) process (fig. S10), which was consistent with decreased forces on nesprin 2 and lamin A in EMT (*20, 38*). Taken together, these results demonstrated the ability of NuTS2 in global dynamic force measurement across SUN2 protein. We also measured the global force changes on SUN1 by using NuTS1, and modest FRET ratio change was observed (fig. S11), implying SUN1 may have less contribution to the force transmission to nucleus in cell spreading compared to SUN2.

### NuTS responds to local force changes with high spatiotemporal resolution

To test the sensitivity of NuTS in response to local mechanical stimulation, we either designed a 5-μm width microfluidic channel to squeeze cells, or used a micropipette to press cells at a local point, as illustrated in Fig. 5A and fig. S12. When suspension HeLa stable cells expressing NuTS2 traveled through microchannels, there was obvious FRET decrease with large deformation occurring at the region that interacted with microchannel (Fig. 5C). When adhesion cells were pressed firmly by micropipette, NuTS2 as well as NuTS1 displayed transient FRET increase in respond to the local mechanical stimulation (Fig. 5D and fig. S13). Once the external forces disappeared, the FRET signal at local region got recovered in both cases (Fig. 5, C and D), demonstrating the excellent sensitivity of NuTS1 and NuTS2 in responding to local tension force changes on SUN proteins with high spatiotemporal resolution. In addition, we reasoned that the high tension exerted on nuclear membrane in both scenarios was by nuclear deformation (*58*).

**Fig. 5.**
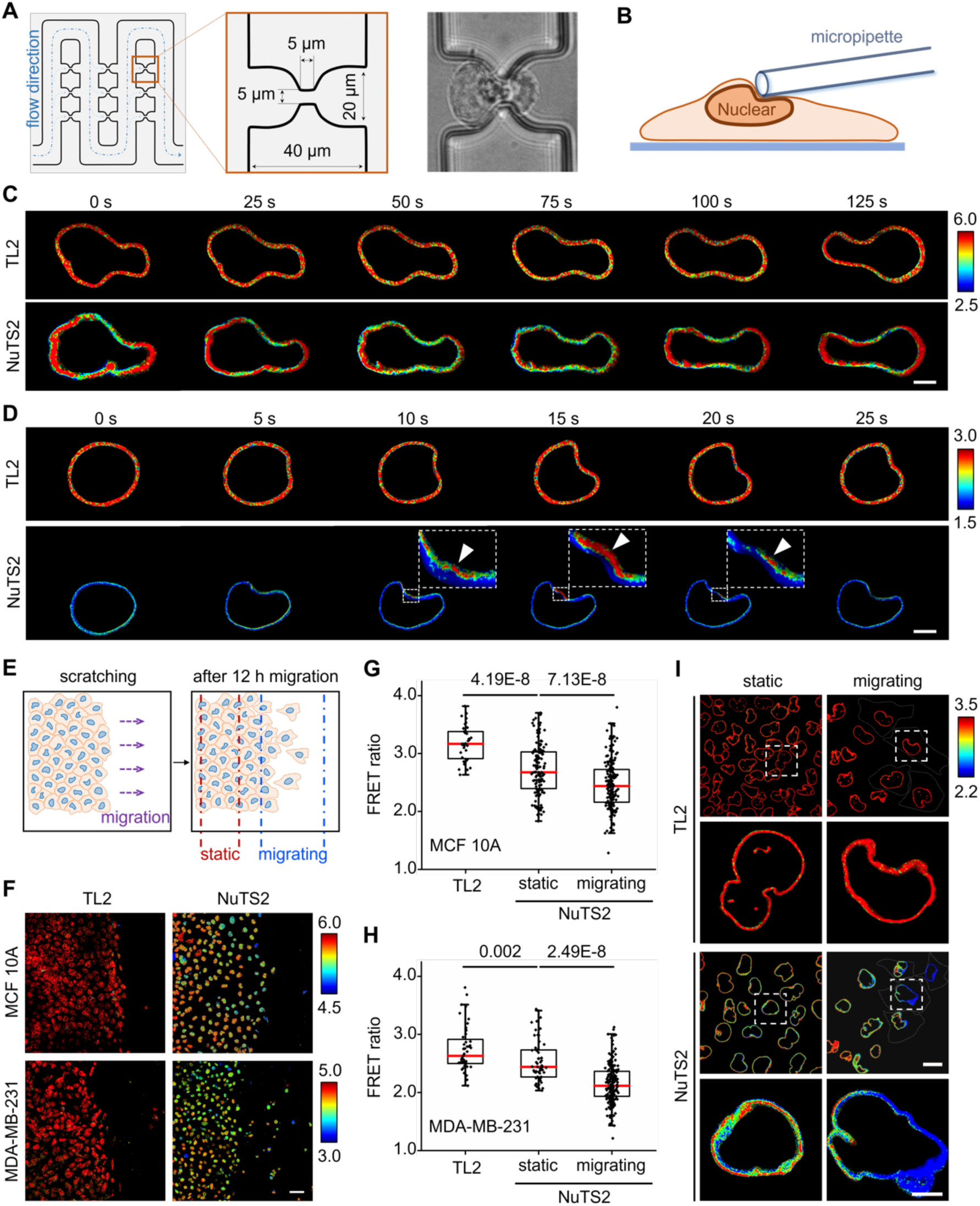
NuTS responds to local force changes with high spatiotemporal resolution. (**A**) The schematic diagram of microfluidic chamber. The middle chart shows the width and height of the flow channel are 40 μm and 30 μm, respectively. The DIC image is a representative image to show cells undergoing squeezing. (**B**) Schematic diagram of pressing the cell with a micropipette in 5 μm diameter. (**C**) Time-lapse imaging showed the dynamic FRET ratio changes when cells were undergoing squeezing. Representative FRET ratiometric images of stable HeLa cells expressing TL2 and NuTS2, respectively. Scale bars, 10 μm. (**D**) The dynamic FRET ratio changes with the continuous pressing of cells expressing NuTS2 by micropipette to generate local mechanical forces. Representative FRET ratiometric images of Hela cells transiently expressing TL2 and NuTS2. Blowup images showed the specific local region with FRET increased. White arrows indicate the local force stimuli. Scale bars, 10 μm. (**E**) The scheme of cell migration in wound scratch assay. The left unscratched areas were defined as “static” areas and the scratched cell-free areas were defined as “migrating” areas. (**F**) Representative FRET ratiometric images of stable MCF 10A cells and MDA-MB-231 cells expressing TL2 and NuTS2 during migration, respectively. Scale bars, 50 μm. (**G**)(**H**) The quantified FRET ratio (Mean ± SEM) from TL2 and NuTS2 in static and migrating stable MCF 10A cells (**G**) and stable MDA-MB-231 cells (**H**). Migrating cells had much lower FRET ratio in both type of cells than static cells. n = 37, 144, 197 (**G**), and n = 48, 61, 203 (**H**). three biological replicates, respectively. Ordinary one-way ANOVA Tukey’s multiple comparisons. (**I**) Representative FRET ratiometric images of stable MCF 10A cells expressing TL2 and NuTS2 during migration. Scale bars, 50 μm for large view and 10 μm for zoom in.

Cell migration is a highly dynamic and polarized process driven by cytoskeleton (*49*). During migration, polarized cytoskeleton regulates the formation of protrusive leading edge in the front and a retracting trailing rear, which transmits different forces into nucleus polarizingly. To visualize nuclear force polarization in living cells, we performed wound scratch assays to generate cell migration model with MCF 10A and MDA-MB-231 cells. 12 hrs post scratching, we observed more migrating cells with MDA-MB-231 cells that had metastatic potential, compared to non-tumorigenic MCF 10A (Fig. 5, E and F). Both types of cells migrated out of the confluency area (static) into cell-free scratching area (migrating) with gradient FRET decrease (Fig. 5, F-H). Strikingly, some population of the migrating cells showed much lower FRET signals in the direction of front leading edge than their retracting trailing rear (Fig. 5I), indicating our NuTS2 can respond to the polarized force transmission on the same nucleus with high spatial resolution. Thus, these results demonstrated that NuTS may provide an efficient imaging tool for monitoring nuclear force changes that respond to normal cell migration or tumor metastasis.

### The nuclear force increases as notochord matures in the zebrafish embryo development

As we demonstrated in living cells, NuTS2 exhibited a rapid and sensitive response to mechanical force changes across SUN2 protein, including actin-mediated cell contractility, LINC complex integrity and extracellular matrix stiffness. Next, we wanted to see how NuTS2 behaves *in vivo*. It is well-known that notochord expands along the direction of the somite growth starting from tailbud during embryonic development, the vacuolation and maturation of the notochord contribute to its elongation and stiffening (*60*). This gives us a hint that the notochord cells must undergo gradient mechanical force stimulation along the anterior notochord maturation direction during embryonic development (Fig. 6A). To verify this hypothesis, we injected NuTS2 mRNAs or TL2 mRNAs into newly born zebrafish embryos with *kdrl*:mCherry genetically modified (*Tg(kdrl:mCherry)*). Kdrl, as a mesoderm marker (*61*), was used to indicate notochord location and structure. FRET imaging *in vivo* was performed when the embryos developed to around 24 hours post-fertilization (hpf) (Fig. 6B). Results showed that NuTS2 was expressed correctly in the whole zebrafish embryo including notochord (fig. S14). The gradient decrease of FRET ratio with NuTS2 was observed from posterior tail buds to the anterior during embryonic development, but without changes in TL2 group (Fig. 6C), which was consistent with the observations in zebrafish embryos expressing lifeAct-mCherry marker (fig. S15). Time-lapse imaging further showed dynamic vacuolation and maturation of notochord cells in the same region of somite from 20 hpf to 23 hpf, vacuoles in notochord cells became bigger while FRET ratio decreased (Fig. 6, D-G), indicating the increasing of nuclear force transmission on SUN2 protein as development progressed in the zebrafish.

**Fig. 6.**
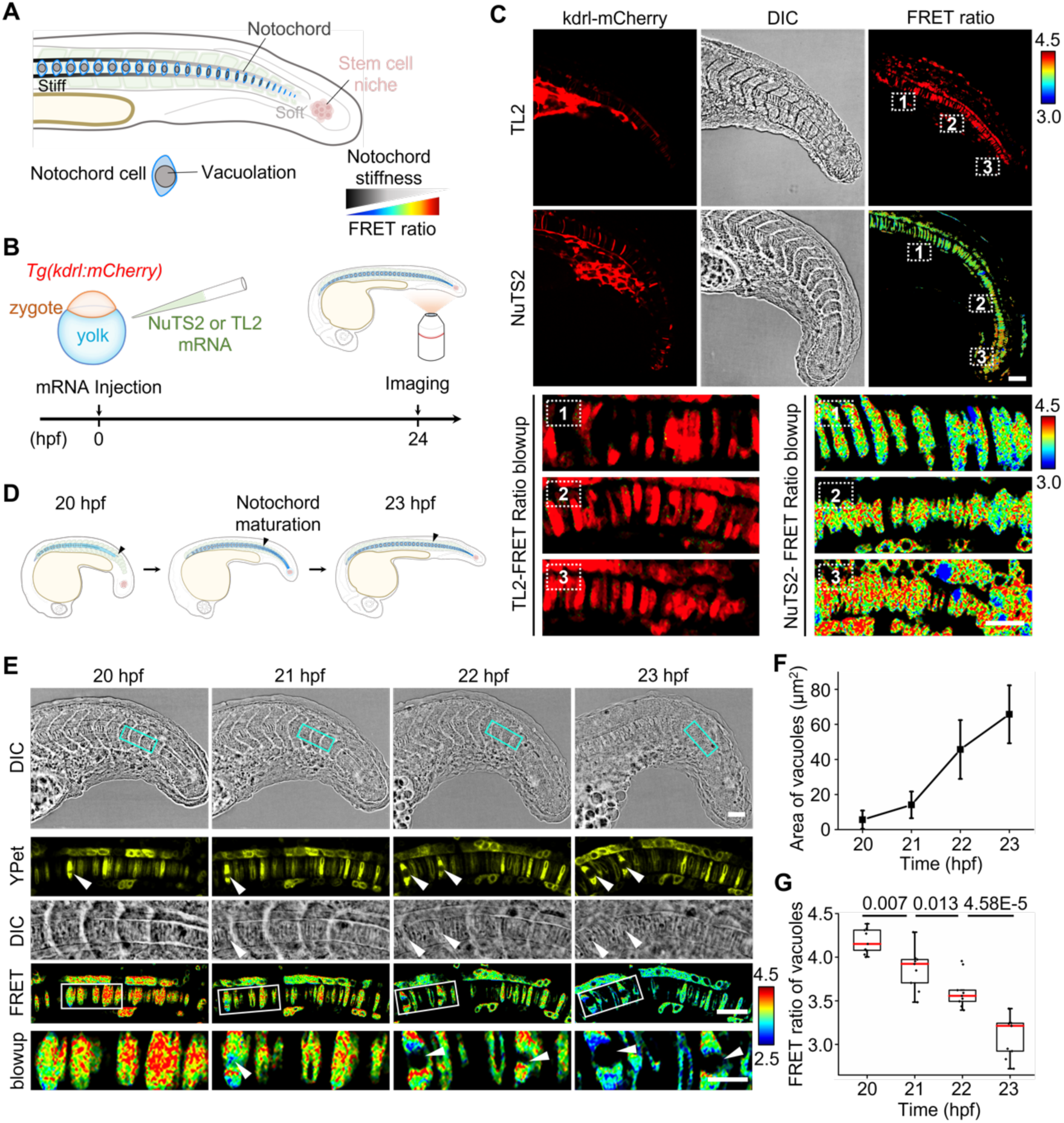
The nuclear force increases across SUN2 protein as notochord matures in the zebrafish development. (**A**) Schematic diagram of somite compartment and notochord. The notochord is vacuolated and expanded during maturation to form a stiff matrix (dark gray), and the soft stem cell niche was located at the end of fishtail (light gray). (**B**) Scheme shows the procedure of mRNA injection into zebrafish embryo with *kdrl*:mCherry genetically modified (*Tg(kdrl:mCherry)*) and imaging process. The mRNA was injected into 1-2 embryoic cells, then imaging was performed on the notochord in fishtail by confocal microscopy. (**C**) Representative FRET ratiometric images in zebrafish embryo with TL2 or NuTS2 expressed, respectively. Kdrl:mCherry was used to indicate the location and structure of notochord. DIC images showed the clear somite compartment. Three regions in FRET ratio image with dashed rectangles were blown up further at the lower panel. FRET ratio decreased from region 3 to region 1 measured by NuTS2. Scale bars, 50 μm for upper images, 20 μm for lower panel images. (**D**) Schematic diagram of notochord vacuolation and expansion during embryo development from 20 hpf to 23 hpf. The black arrow shows that the same location of notochord was imaged during its maturation. (**E**) The DIC images represent developmental process of body segments at 20, 21, 22, and 23 hpf, respectively. Scale bars, 50 μm. Representative FRET ratiometric images at the same region of somite (the blue rectangular boxes) during notochord maturation were blown up. White arrows show the vacuolation of notochord. Scale bars, 20 μm. (**F**) The curves of vacuoles areas at different developmental time points from (**E**). The areas of vacuoles were increased during notochord maturation. n = 3. (**G**) The quantified FRET ratio (Mean ± SEM) of NuTS2 from (**E**). n = 9. Ordinary one-way ANOVA Tukey’s multiple comparisons.

### Forces on SUN2 protein are required for zebrafish embryo development

To investigate the correlation between nuclear forces and notochord maturation, on one side, we used Bafilomycin A1 (Baf A1), an H+-ATPase inhibitor, to inhibit lysosomal acidification in embryonic notochordal cells, thus block notochordal vacuolation and maturation (*60*). On the other side, we used Cyto D, which inhibits actin polymerization and has been used in live cell imaging in Fig 2, to serve as a positive control to reduce cell contractility. Zebrafish embryos were treated with drugs for 4 hrs starting from 20 hpf (Fig. 7, A and B). At 24 hpf, DMSO group showed big vacuoles in notochord cells near cloaca in both YPet and mCherry channels, but there were no vacuoles at all in Baf A1 group, or tiny ones in Cyto D group, which suggested that drugs had effects on notochord vacuolation and maturation (Fig. 7C). Then, we looked at nuclear force changes by NuTS2. Notably, FRET ratio increased significantly in the drug treated groups, particularly Cyto D treatment increased FRET signal up to two-folds compared to DMSO group (Fig. 7, C and D), indicating nuclear forces across SUN2 protein was reduced by both Baf A1 and Cyto D. To see if there are any functionality deficiency caused by the reduction of nuclear force transmission, we measured tailbud bending angle (*Ɵ*), number of somites at tailbud (*N*), as well as the length of 5-somite (*l*) near cloaca in DMSO group and drug treatment groups during tailbud stages of development (Fig. 7E). Our quantitative results showed that number of somites at tailbud (*N*) was significantly decreased by both Baf A1 and Cyto D (Fig. 7, F and G). N at 20 hpf was 3.75±1.16 in average, after 4 hrs, DMSO group increased to 10.67 ±0.51, Baf A1 group was 4.86±0.90, and Cyto D group was 8.50±1.05 (Fig. 7G). Regarding changes in tailbud bending angle (*Ɵ*) and length of 5-somite (*l*), they were only significantly affected by Baf A1 but not Cyto D (Fig. 7, H and I). This possibly implied that there were more mechanical forces abolished in notochord cells by inhibiting vacuolation than cytoskeleton contractility. Putting together, vacuolation does generate more forces in notochord cells and transmit tension force from cytoskeleton into nucleus through SUN2 protein as notochord expands and matures. Disrupting either vacuolation or force transmission affects notochord maturation and zebrafish embryo development.

**Fig. 7.**
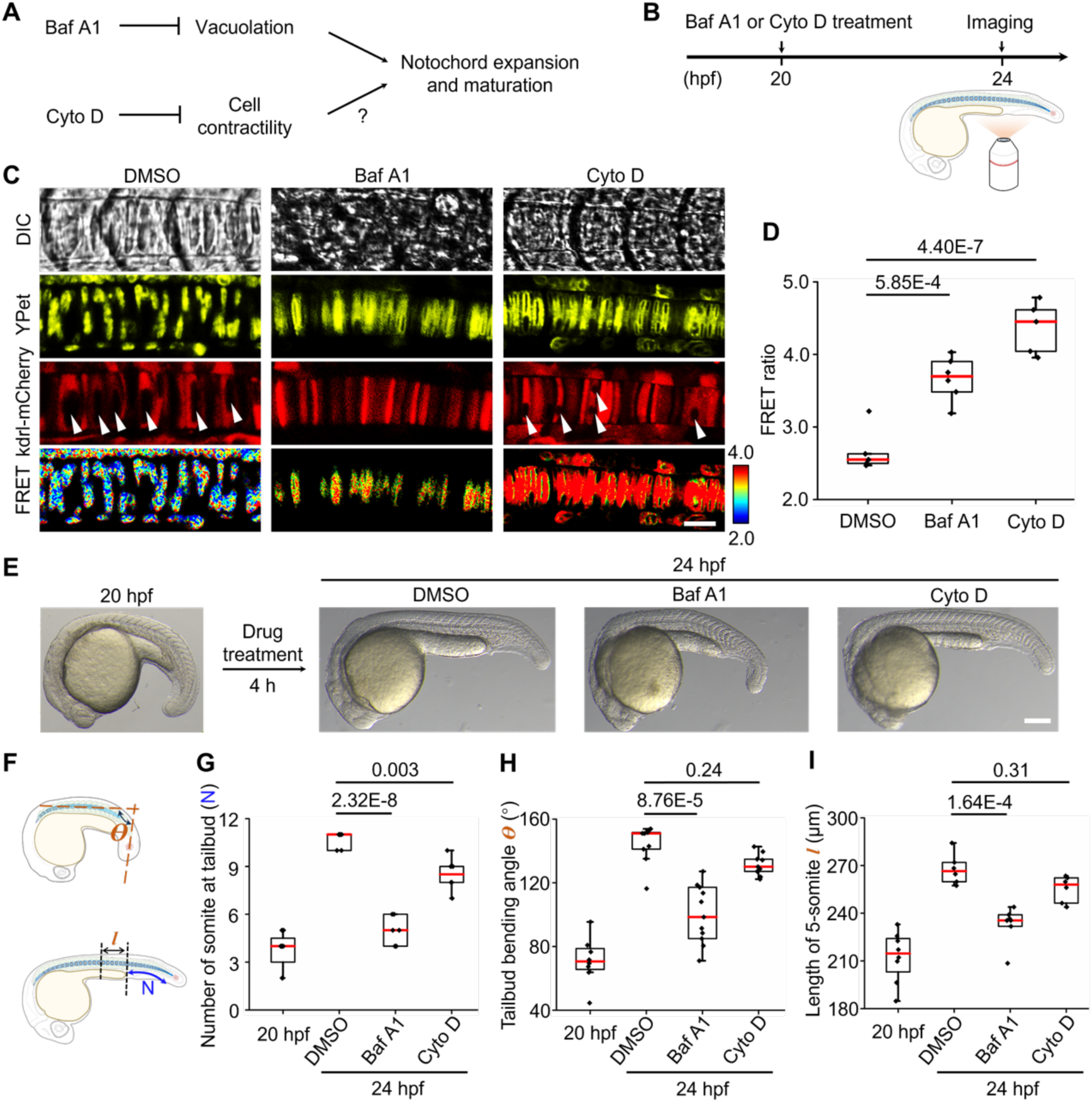
Nuclear forces on SUN protein affect notochord maturation and zebrafish embryos development. (**A**) The scheme of drug treatment for manipulating notochord expansion and maturation. Baf A1 effects notochord vacuolation while Cyto D reduces cell contractility. (**B**) Schematic diagram of drug treatment and imaging procedure. The 20 hpf embryos were treated with 0.5 μM Baf A1 and 1 μM Cyto D for 4 hrs, respectively, and then the imaging data were collected at 24 hpf. (**C**) Representative FRET ratiometric images of notochord maturation with different drug treatment. White arrows show the vacuolation of notochord. Scale bars, 20 μm. (**D**) The quantification of FRET ratio (Mean ± SEM). FRET signals increased dramatically with both Baf A1 and Cyto D treatment compared to DMSO group. n = 5. (**E**) Effects of Baf A1 and Cyto D treatments on the developmental defects in terms of embryos phenotype. Representative DIC images of notochord maturation with different body somite extension and bending angle. Scale bars, 100 μm. (**F**) The schemes showed the definition of tailbud bending angle, length of 5-somite, and number of somites at tailbud. *Ɵ* represents the angle between the parallel line of the body segments of the fish belly (dashed horizonal line) and the parallel line of the fish tail (dashed vertical line). *l* represents five-somite region before the cloaca. *N* represents number of somites at tailbud (from cloaca to tail). (**G**) The quantification of somite number (N) at tailbud (Mean ± SEM). n = 8, 6, 7, 6. (**H**) The quantification of bending angle *Ɵ* (Mean ± SEM). n = 8, 9, 11, 9. (**I**) The quantification of length of a five-somite region (*l*) (Mean ± SEM). n = 8, 6, 7, 6. Ordinary one-way ANOVA Tukey’s multiple comparisons.

## Discussion

Despite notable advances, the dynamics of force transmission from the cytoskeleton to the inner nuclear membrane via the LINC complex remains largely unexplored. Here, we developed FRET-based nuclear tension sensors, NuTS1 and NuTS2, to measure tension forces on SUN1 and SUN2 proteins in living cells and zebrafish embryos. We discovered that both the coiled coil structure and SUN domains of the SUN protein were required for NuTS to sensitively respond to force changes originating from the cytoskeleton. Our findings demonstrated the potential of NuTS1 and NuTS2 to measure forces and study the effect of force dynamics across the nuclear membrane. Notably, NuTS2 can rapidly and sensitively respond to global and local mechanical stimuli *in vitro* and *in vivo* with high spatiotemporal resolution, including actin-mediated cell contractility, extracellular matrix stiffness, chromatin decondensation, EMT transition, cell spreading, cell migration, embryo development, and so on. This highlights the importance of SUN2 in force transmission through the LINC complex in nuclear mechanosensing and mechanoadaptation.

The mechanical force on the outer nuclear membrane has been measured by a number of nesprin tension sensors that have been developed previously (*20, 21, 34, 62*). Their approaches were quite similar, by employing the general tension sensor module (TSMod) (*32*) into the mini Nesprin-2G protein (*63*). Mini Nesprin-2G contains N-teminal calponin homology domains CH1 and CH2 for actin binding, C-terminal transmembrane KASH domain for SUN protein binding, and spectrin repeats SR1, SR2, SR55, and SR56 in the middle for TSMod placement. There were two ways to place TSMod, one was in between SR2 and SR55 (*34, 62*), and the other was after SR56 (*20, 21*). No matter where TSMod was inserted into mini nesprin-2G, there were large number of spectrin repeats (SR3-54) missing that included the interface to engage microtubules and microtubule motors (*64*), resulting the role of microtubule forces cannot be addressed by previous nesprin tension sensors. Distinguishingly, our SUN tension sensors connect to the intact nesprin proteins in endogenous, and measure all the tension forces from cytoskeleton proteins to inner nuclear membrane through nesprin, therefore, the information provided by our NuTS is substantially different from that of the previous studies reporting tension levels on a truncated nesprin. Moreover, SUN forms trimer and its C terminus connects to nesprin proteins through KASH lid domain (*41*). The force level measured using NuTS is essential for understanding the mechanical stability and responses of SUN complexes, as well as their connections to lamin A network at the inner nuclear membrane. Therefore, the insights gained from our measurement are distinctive and substantial, which cannot be derived from previous studies.

NuTS not only provides imaging tools for direct force measurement across SUN proteins but also serves as an indicator of nuclear force changes under various physiological and pathologic processes regulated by mechanical forces, such as embryo development (*60*), tumor metastasis (*11*), aging (*65*), arteriosclerosis (*4*). Emerging evidence suggests that the nucleus is a crucial mechanical sensor and effector (*28*). Mechanical forces from extracellular microenvironment can be transmitted through the pathway of “focal adhesion-cytoskeleton-LINC complex” pathway to the nucleoskeleton and nucleus interior (*27*), triggering nuclear deformation, chromatin redistribution and gene expression (*5, 17*). SUN proteins, key structural components of the LINC complex, play a vital role in transmitting forces into nucleoskeleton proteins like lamin A. In our study, both NuTS1 and NuTS2, which incorporate with F40 linker, exhibited notable FRET changes when small molecules disrupted actin-mediated cell contractility (Fig. 2), indicating that NuTS senses force changes through the actin-LINC complex pathway. The F40 linker behaves like a molecular spring with gradual elongation under forces between 1 to 6 pN (*32*). Although both nesprin-2G (*34*) and our NuTS measured different forces on nesprin and SUN proteins via the F40 linker, further investigation using single-domain peptides for biosensor optimization is necessary for more accurate measurement of the gradient force transmission across LINC complex.

Forces across SUN proteins are transmitted radially within the nuclear membrane, differing from the tangential tension force on the nuclear lamina measured by an intermolecular tension FRET biosensor based on nanobodies (*38*). The specificity of NuTS was validated in our study by either nesprin knockout or lamin A knockout (Fig. 3), confirming that it measures force transmission from the cytoskeleton to nucleus through an intact LINC complex as previously reported (*38*). Our study also revealed that NuTS2, but not NuTS1, exhibited greater sensitivity to the connection between nesprin 2 and lamin A, suggesting that SUN2 plays a major role in nuclear force transmission. This finding aligns with reports that the actin, LINC complex (nesprin and SUN2), and lamina compose nuclear lines providing mechanical linkage between cytoskeleton and nucleoskeleton in stretch stimulation (*66*). To explain why there was no changes on NuTS1 in nesprin 2 knock out cells (fig. S4, B and C), it is likely due to the requirement of specialized SUN: nesprin pair for their function (*48*), which means SUN1 primarily interacts with nesprin 1 but not nesprin 2, in the force transmission. In addition, the downstream connections of SUN1 have several possible nuclear membrane proteins, including lamin A (*29*), Nup153 (*49, 50*), and emerin (*51*), to not only transmit forces but also play roles in stabilizing nuclear pore complex and association with other inner nuclear proteins, which could be the alternative explanation for NuTS1 not sensitive to *LMNA* depletion. *LMNA* depletion may not sufficiently affect the state of SUN1.

Nuclear deformation remarkably contributes to the regulation of cellular functions such as cell migration and pathogenesis (*58*). Utilizing NuTS2, we observed a heterogeneous force transmission into the nucleus when suspended HeLa cells underwent large deformation in the microfluidic channel. The anterior and posterior regions of the nucleus exhibited much lower tension forces compared to the squeezing area (Fig. 5, A and C), suggesting that severe heterogeneous mechanical force could lead to localized loss of nuclear envelop integrity, chromatin herniation, DNA damage, and micronuclei formation ((*18, 25, 26*). Interestingly, when single-point stress was applied to adhering HeLa cells by micropipette pressing, less force was transmitted on NuTS2 across the nuclear envelop at the poking area (Fig. 5, B and D), showing different responses compared to cell squeezing. We speculated that difference in cytoskeleton structures corresponding to different cell status primarily affect nuclear force transmission. Stress fibers are absent in suspended cells but present for force transmission in adherent cells (*67*). Additionally, collective cell migration in 2D cultures is often used to simulate developmental and pathological processes, such as wound healing. Migrating leader cells crawl out from the cluster sheet with polarized cytoskeleton (*68*), and the polarized cytoskeleton pulling the cytosol and nucleus forward (*69*). Thus, the nuclear forces may be enhanced when the pulling force transmit to nuclear envelop. NuTS2 demonstrated that migrating leader cells show higher tension forces on SUN2 protein than the static follower cells (Fig. 5, E-I), providing visual and intuitive evidence of nuclear mechanics during cell migration.

Furthermore, NuTS offers an excellent imaging method for nuclear force measurement and its regulatory function investigation *in vivo*. During development, tissue formation and maturation are usually accompanied by changes in biophysical property such as tissue stiffness (*70, 71*). These mechanical forces are transmitted into the cells as well as nucleus to determine cell behavior and fate (*7*). It has been reported that the tail buds of zebrafish embryos, the center of somite progenitor cells, have low tissue stiffness, which increases along the anterior direction of the somite (*72*). Recent studies suggest that the notochord cells derived from tailbuds with vacuolation and maturation during development, which provides an internal expansion stress to contribute to notochord elongation and stiffening (*60, 73*). Both pressure and stiffness could affect the magnitude of tension force transmission from cytoskeleton to nucleus, through anchoring points (LINC complex) on nuclear membrane, including tension force transmission that SUN2 protein anticipates. Our studies observed that the nuclear mechanical forces were elevated with the tissue maturation, and the dynamic force changes can be directly visualized and measured by NuTS system. Disrupting either vacuolation or actin polymerization causes decrease of tension force transmission on NuTS2 significantly (Fig. 7). Therefore, NuTS2 measures tension force changes in the notochord which relates to actin-induced forces in mammalian cells. This biomechanical cues in nucleus are critical for embryo development.

In summary, NuTS enables the direct imaging of nuclear force transmission across SUN proteins with high sensitivity and high spatiotemporal force resolution in response to mechanical stimuli both *in vitro* and *in vivo*. Our findings provide important insight into the dominant role of SUN2, rather than SUN1, in the force transmission of nuclear mechanics across LINC complex, encompassing cellular processes and embryo development. This work opens up the possibility of utilizing FRET-based nuclear tension sensors to measure nuclear forces and elucidate its regulatory mechanisms in known and unknown physiological and pathologic processes.

## Materials and Methods

### Designation and construction of the SUN tension sensors (TS)

To build a series Förster resonance energy transfer (FRET) -based TSs, the mouse NuTS1 (for SUN1) and NuTS2 (for SUN2) were designed based on the previous method of force measurement propagation across target proteins (*40, 74*). In brief, SUN1 and SUN2 expression constructs are based on murine Mus musculus Sad1 and UNC84 domain containing 1 (SUN1) and Mus musculus Sad1 and UNC84 domain containing 2 (SUN2) cDNA (NCBI Reference Sequence: NM_024451.2 for SUN1 and NM_001205345.1 for SUN2). The tension linker F40, containing 40 amino acids long flagelliform peptide (GPGGA)8 (*32*), was used to chemically synthesize with two NheI sites in pUC-GW-Amp vector by GENEWIZ® from Azenta Life Sciences. Next, the F40 was obtained from digest product by using NheI restriction endonuclease (New England Biolabs, cat #R3131S), and they were insert between FRET pairs-ECFP and YPet in a pSIN-Amp+ vector by using T4 ligase (New England Biolabs, cat #M0202S), and series ECFP-tension linker-YPet segments were prepared by PCR. To examine the effect of the TS and optimize its size, the subunit LMNA-binding domain (1-139 aa for NuTS1 and 1-128 aa for NuTS2) and transmembrane domain (TM, 416-455 aa for NuTS1 and 227-266 aa for NuTS2) that connected by a GGSGGT linker were selected for all TS. The KASH lid segment for KL-NuTS (737-781 aa for KL-NuTS1 and 555-599 aa for KL-NuTS2), SUN domain for SD-NuTS (737-913 aa for SD-NuTS1 and 555-731 aa for SD-NuTS2), and CC-SUN for CSD-NuTS (456-913 aa for CSD-NuTS1 and 267-731 aa for CSD-NuTS2) were prepared by PCR, respectively. The zero-force FRET control in the absence of tension was constructed by removing the C-terminate tail behind FRET pairs (called tailless, TL). The main primer sequences used are listed in table S1. Finally, all NuTSs were constructed with Gibson Assembly system by using GeneArt™ Gibson Assembly HiFi Kit (Invitrogen, Thermo Fisher Scientific, cat #A46628).

### Cell culture

HeLa cells was cultured in high glucose Dulbecco’s Modified Eagles Medium (DMEM) (Gibco, Thermo Fisher Scientific, cat #C11995500BT) containing 10% (vol/vol) fetal bovine serum (Gibco, Thermo Fisher Scientific, cat #10099-141C), 100 units/mL penicillin-streptomycin (Thermo Fisher Scientific, cat #15140122). C2C12 was cultured in high glucose DMEM containing 20% (vol/vol) fetal bovine serum, 100 units/mL penicillin, and 100 units/mL streptomycin. For transfection of NuTS/TL, the cells were seeded in 24 wells plate for 12 h, HeLa cells was transfected with 500 ng plasmid (1 μg for C2C12) by Lipofectamine™ 3000 Transfection Reagent (Invitrogen, Thermo Fisher Scientific, cat #L3000001) for 24 h, then transfer to glass bottom dishes for 12 h and imaged. All cells were cultured in incubator at 37°C with a 5% CO_2_ in humid air atmosphere.

### Drug treatments

The target cells were transfected with NuTS and TL plasmid for 24 h, respectively, then transferred to glass bottom dishes for another 12 h culture for drug treatment. To decrease the mechanical force transmitted from cytoskeleton, cells were treated with Cytochalasin D (MedChemExpress, KLink Biotechnologies, cat #HY-N6682) at a concentration of 1 μM for 0.5 h, or treated with (-)-Blebbistatin (Sigma-Aldrich, cat #203391) at a concentration of 30 μM/mL for 1 h, respectively. To inhibit Rho A kinase to reduce myosin activity, cells were treated with Y-27632 (Cell Signaling Technology, cat #13624S) at a concentration of 10 μM for 1 h. To increase cell contractility, cells were treated with Calyculin A (Cell Signaling Technology, cat #9902S) at a concentration of 1 nM for 30 min. For EMT induction, cells were treated with recombinant human TGF-beta1 (Cell Signaling Technology, cat #75362) at a concentration of 5 ng/mL for 24 h and 48 h. To decondense chromatin, cells were treated with trichostatin A (TSA) (Cell Signaling Technology, cat #9950S) at a concentration of 500 nM for 4 h to increase euchromatin. All controls were treated with 1‰ DMSO for 1 h. The images for dynamic FRET ratio changes under Cyto D or Y-27632 treatment were taken per 15 min. To disrupt LINC complexes, the dominant negative KASH (DN-KASH)-mCherry plasmid was transfected into the TL and NuTS stable HeLa cells, the images were taken after 36-48 h transfection.

### FRET imaging acquisition and data analysis

The approach we used was the intensity-based ratiometric FRET (fig. S16). Firstly, the samples were imaged using Dragonfly spinning disk confocal microscopy (Seven-laser, 60× Oil (NA1.45)) (Dragonfly CR-DFLY-202-2540, U.S.A.), equipped with a cooled charge-coupled device (CCD) camera, a 445DF46 excitation filter, a 450DRLP dichroic mirror and two emission filters controlled by a filter changer (478DF37 for ECFP and 521DF38 for FRET), to collect fluorescence data by Andor Fusion 2.3.0.44, and then Imaris 9.9.0 software was used to export fluorescence intensity images of ECFP, YPet and FRET channels as .tiff files, which can be recognized by our FRET imaging analysis software Fluocell in the next step. Subsequently, according to our previous method (*75*), the image analysis software package Fluocell based on MATLAB (http://github.com/lu6007/) was utilized to generate FRET ratio data and images from each cell. FRET ratio in live cells was calculated just on the nuclear envelop rim by Fluocell, in which there is a customized program for detecting the obvious rim with high intensity and measuring average FRET ratio in the rim region only. The path under Fluocell to find the customized program is: fluocell-master/app/quantify/quantify_ratio_intensity. Since the TL contains only the N-terminal and transmembrane peptides of the SUN proteins, there is some amount of tension sensors diffused in the cytoplasm, so does KL-NuTS and SD-NuTS due to trunctated C-terminus. In order to highlight FRET images on the nuclear membrane, we used the similar strategy for our image display by following Nesprin TS papers in 2016 Biophysics (*34*) and Gomes & Borghi 2020 (*20*). In brief, we selected the same representative cell from both YFP channel and FRET channel to get zoomed in (“duplicate” in Image J/Fiji), and then merged the YPet channel indicating the localization of NuTS and FRET channel by using the merge channel function of Image J/Fiji. Since there was almost no fluorescent signal inside nucleus, we can easily tell where the inner nuclear membrane was. With the merged images, based on the localization of YPet on nuclear membrane, the outer nuclear boundary can be identified and circled out. Then the nuclear envelop rim was segmented out by “clear outside” function. The FRET ratio in zebrafish was also calculated by Fluocell but with regular steps as what we have done previously (*75*). The average FRET ratio was measured from the entire fluorescent regions but not in the nuclear envelop rim due to the resolution in zebrafish.

### Statistical analysis

Experiments were performed in triplicates at least unless otherwise stated. Quantitative values were expressed as the Mean ± SEM. The Ordinary One-way Analysis of Variance (ANOVA) test technique followed by a Tukey’s post-hoc test with Origin Pro 2016 was utilized to assess the level of statistical significance.

## Acknowledgments

We thank Dr. Xiaohu Zhou, Chang Sun, and Fei He at Shenzhen Bay Laboratory for discussing microfluidic chamber design and helping with SUN KO cells and micropippette experiments, respectively, and Dr. Wenhua Yan at The Second Affiliated Hospital of Chongqing Medical University for providing the *Tg(kdrl:mCherry)* zebrafish. We thank the Bioimaging Core facility at Shenzhen Bay Laboratory for providing imaging support. We also would like to acknowledge engineers Mei Yu and Shixian Huang from the Bioimaging Core facility for assistance with Dragonfly Spinning Disk Confocal Microscopy (7-laser).

## Funding

National Key R&D Program of China (2022YFC2704303 to J.Z. and 2022YFF0710700 to J.H.Q.)

National Science Foundation of China (32100450 to Q.P. and 32325030, 82270419, and 81921001 to J.Z. and 12372302 to J.H.Q.)

Shenzhen Medical Research Fund (B2402010 and D2301009) Guang-dong Pearl River Talent Program (2021QN02Y781)

Shenzhen Bay Scholars Program

The Open Sharing Fund for the Large Instruments and Equipments of Shenzhen Bay Laboratory

## Author contributions

Conceptualization: JQ, JZ, QP Methodology: JW, SL Software: SL

Validation: JW, SD, FX Formal analysis: JW, Investigation: JW, SD, YL Resources: JW, FX

Data curation: JW,

Writing-original draft: JW, JQ, QP Writing-review & editing: JW, JQ, JZ, QP Visualization: JW, SL, QP

Supervision: YL, JQ, JZ, QP Project administration: QP Funding acquisition: JZ, QP

## Data and materials availability

All data are available in the main text or the supplementary materials.

## Supplementary Materials

Materials and Methods Table. S1.

Figs. S1 to S16 Movie S1 to S2 References (*60*)

## Supplementary Text

### Materials and Methods

#### Establishment of Knockout cell lines with CRISPR/Cas9

For *SYNE2* KO C2C12, the single guide RNAs (sgRNAs) were designed against *SYNE2* gene N-terminus in exon 2 genome (https://www.ncbi.nlm.nih.gov/nuccore/NC_000078.7, GeneID: 319565) with an online guide design tool (https://cctop.cos.uni-heidelberg.de:8043/). Two pairs of sg RNA (spacer I, spacer II) were designed, *SYNE2* target sequence 1: GAAACCCTCTCCATCTTCCGTGG, and target sequence 2: GCGGCCATGCTTTGACTCAGTGG, the forward primer 5’-CACCGAAACCCTCTCCATCTTCCG-3’ and reverse primer 5’-AAACCGGAAGATGGAGAGGGTTTC-3’ for target sequence 1 (T1), the forward primer 5’-CACCGCGGCCATGCTTTGACTCAG-3’ and reverse primer 5’-AAACCTGAGTCAAAGCATGGCCGC-3’ for target sequence 2 (T2) were synthesized by GENEWIZ® from Azenta Life Sciences. The forward and reverse primer was programmed annealing for each target sequence. Next, BbsI endonuclease cleaved pX458 vector was connected with target sequence by using T4 ligase. For expression, the sgRNA_*SYNE2*_T1 (500 ng) and sgRNA_*SYNE2*_T2 (500 ng) in pX458-SV40 NLS-Cas9-ECFP) were transfected by using the Lipofectamine 3000 transfection reagents (Invitrogen, cat #L3000015) followed by selection of ECFP-positive cells with Cell Sorter (CytoFLEX SRT, Beckman). The KO cells was verified by Immunofluorescence staining and further sequenced to specifically detect the knockout region in gene. Same approach was applied for generating the stable cell lines of *SUN1*, *SUN2*, and *LMNA* KO C2C12.

#### Immunofluorescence staining

For identify the stable KO cell lines, cells were seeded at a density of 10,000 cells/cm^2^ in 10% serum medium for 12 h, rinsed with PBS for 3 times, then fixed in 4% (w/v) paraformaldehyde for 30 min and rinsed with PBS for 3 times, permeabilized in 0.25% (v/v) Triton X-100 for 10 min and rinsed with PBS for 3 times. Subsequently, the non-specific binding sites were blocked by incubating with 5% BSA for 2 h at room temperature. The primary antibody, Anti-SUN1 antibody (1:200; abcam, cat #ab103021) for SUN1, Anti-SUN2 antibody (1:500; abcam, cat #ab124916) for SUN2, Syne-2 Antibody (F-11) (1:200; Santa Cruz, cat #sc-398616) for nesprin 2, lamin A/C (4C11) Mouse mAb (1:200; Cell Signaling Technology, cat #4777S) for lamin A/C, respectively, was overnight incubation at 4°C and rinsed with PBST (PBS with 0.1% Tween® 20). Next, the samples were incubated with secondary antibody at room temperature for 2 h, goat anti-mouse Alexa Fluor® 488 (1:1000; Abcam, cat #ab150077) for SUN1, SUN2 and nesprin 2, and anti-mouse Alexa Fluor® 647 (1:1000; Cell Signaling Technology, cat #4410S) for lamin A/C, respectively. Finally, for F-actin staining, the cells were incubated with Rhodamine-Phalloidin Alexa Fluor™ 568 (1:1000; Invitrogen, cat #A12380) overnight at 4°C, rinsed with PBST and incubated with Hoechst 33342 (1:2000; Cell Signaling Technology, cat #4082S) for 10 min at room temperature for nuclei staining. After being rinsed with PBST, the samples were mounted in VECTASHIELD Antifade Mounting Medium (Vectorlabs, cat #H-1000-10) and took image by Dragonfly Confocal Microscopy System (Dragonfly CR-DFLY-202-40, U.S.A.).

#### Hydrogel preparation for extracellular mechanics and cell seeding

The Polyacrylamide (PAA) hydrogel on glass bottom dish was prepared as **Fig. S5**. Briefly, the 35 mm Petri Dishes with 18 mm well (0# Germany Cover glass, Cell E&G) were following treated with 0.1 M NaOH (Sangon Biotech) for 40 min, 3-Aminopropyltrimethylsilane (Sigma-Aldrich, cat #440140) for 5 min, washed with ddH_2_O for 3 times, covered with 0.5% glutaric dialdehyde (Sangon Biotech) for 30 min, washed with ddH_2_O for 3 times. Next, the polyacrylamide hydrogel premix of 0.2 kPa and 40 kPa were dropped and cover with coverslip for flatten, respectively, the coverslip was peeled off in ddH_2_O after curing for 30 min. Then, the gel was immersed into the crosslinker of 0.25 mg/mL Sulfo-SANPAH (Thermo Fisher Scientific, cat #22589) and exposed in UV for 10 min, washed with PBS for 3 times. Finally, the gel was coated with 20 μg/mL fibronectin overnight at 4°C and 0.1% gelatin for 1 h at 37°C. The transfected C2C12 cells were seeded at the density of 10,000 cells/cm^2^, and the images were taken by Dragonfly Confocal Microscopy System (Dragonfly CR-DFLY-202-40, U.S.A.) after 12 h culturing.

#### Cell adhesion assay

The C2C12 cells were transfected with NuTS and TL plasmid for 24 h, respectively, and then cells were digested with 0.05% trypsin and suspended with DMEM containing 20% (vol/vol) fetal bovine serum, 100 units/mL penicillin, and 100 units/mL streptomycin. 1 mL cell suspension with 0.05 million/mL density was added into a glass bottom dish, and the images were taken after 20 min initial adhesion. The images were collected for 3 h, and interval was 10 min/frame.

#### Micropipette pressing assay

The micropipette was prepared by MODEL P-1000 flaming/brown micropipette puller (SUTTER Instrument, U.S.A.). Detailly, Capillary tube (length: 100 mm, outer diameter: 1.0 mm, inner diameter: 0.8 mm) was fixed on instrument, and the following parameters were used: Heat 525, Pull 70, Velocity 70, Delay 50, Pressure 300, the micropipette with

∼5 μm diameter was obtained. Then, the HeLa cells cell transfected with NuTS and TL plasmids for 24 h and seeded into a homemade PDMS U-shaped chamber for another 12 h. Finally, the micropipette was fixed on a microscope stage attached to the Dragonfly Confocal Microscopy System (Seven-laser, 100×Oil (NA1.45)) (Dragonfly CR-DFLY-202-2540, U.S.A.), and the tip of the needle was slowly lowered close to the nucleus area and stimulated with compression. The FRET images were collected at the same time.

#### Generation of stable cell lines

To establish HeLa, MCF 10A, and MDA-MB-231 stable cell lines that constitutively express NuTS2 or TL2, the lentivirus was constructed to infect the cells. In brief, the HEK 293T cells were seeded in 6 wells plate and cultured in standard DMEM with 10% Fetal Bovine Serum, 100 units/mL penicillin, and 100 units/mL streptomycin in a humidified atmosphere of 5% CO_2_ at 37°C. For lentivirus package, the pSIN-TL2 or NuTS2, pCMV-ΔR, and pCMV-VSVG plasmids were co-transfected into HEK 293T cells with 60∼70% confluency using Lipofectamine 3000 transfection reagents (Invitrogen), respectively. The medium was changed with fresh medium after 6 h transfection, and the supernatant was harvested after 48 h, then filtered with a 0.45 μm filter and precipitated by PEG-*it*^TM^ Virus Precipitation Solution (5×, System Biosciences, cat #LV810A-1) at 4°C overnight, centrifugation to concentrate at 4°C 1500 g for 30 min, resuspended with PBS and stored at −80°C. Note, all operations were performed on an ice bath. Next, the virus was applied to infect target cells, and the fluorescence positive cells expressing the tension sensor were sorted by using cell sorter (FACSAria Fusion, U.S.A.).

#### Squeezing assay

The entire preparation includes two processes: fabrication of silicon micropattern masters, and preparation of PDMS micropattern. The fabrication of silicon micropattern masters were entrusted to BYMICROFAB (Suzhou, Jiangsu, China). The silicon masters used in our work contained a micropattern as show in Fig. 4A. Then, PDMS micropattern was generated by replica molding showed in fig. S12A. Briefly, poly-dimethylsiloxane (PDMS, Dow Corning, Sylgard 184) prepolymer with a curing ratio 10:1 was poured over the silicon micropattern masters mold to make a negative template, degassing under vacuum for 30 min and cured at 60°C overnight. Next, the negative PDMS mold was peeled off, oxidized with O_2_ plasma (Plasma PT-5S, Sanheboda, Shenzhen, China) and passivated with trichloro (1H, 1H, 2H, 2H-perfluorooctyl) silane (Sigma-Aldrich, cat #448931) vapor overnight under vacuum. For positive PDMS micropattern preparation, PDMS prepolymer was poured over the PDMS negative template, degassing under vacuum for 30 min, curing at 60°C overnight, and gently peeling off the PDMS micropattern from the negative PDMS template. The final micropattern and a clean glass coverslip was treated with O_2_ plasma for 90 s, 120 W, and then both treated surfaces were rapidly stuck together and bonded. Finally, the stable cell lines of NuTS2 or TL2 were detached from dishes and filtered, and the cell suspension was flowed through the micropattern by using a microinjector (Fusion 200-X, Chemyx), the rate of flow is 0.002 ml/min.

#### Preparation and injection of mRNAs

For mRNA preparation, the plasmids with T7 promoter and target inserts (NuTS2, TL2, and lifeAct-mCherry) were transcribed *in vitro* by using the AM1344 mMESSAGE mMACHINE™ T7 Transcription Kit (Invitrogen, Thermo Fisher Scientific, cat #AM1344). Then, the transcription products were collected by the E.Z.N.A. Total RNA Kit I (Omega, cat #R6834-01), and concentrations were generally in the range of 300-500 ng/μL. For mRNA injection, the micropipettes for microinjection were prepared as described in the micropipette pressing assay. Opening the tip of the needle with a razor blade, and the mRNA of TL and NuTS2 only was injected for *kdrl*:mCherry transgenic zebrafish that gifted by Wenhua YAN (The Second Affiliated Hospital of Chongqing Medical University), respectively. In addition, the mRNA mixture of NuTS2 or TL2 with lifeAct-mCherry were injected 1-2 nL into the embryos at 1-2 cell stage (within 0.75 hours post-fertilization, hpf) of WT zebrafish (AB). Finally, the injected zebrafish embryos were cultured to 18-24 hpf at 28°C for the following experiments.

#### Drug treatment on zebrafish embryos

The chorionic villi of injected zebrafish embryos were peeled off at 20 hpf, and divided equally into three groups. One group was treated with 0.5 μM Bafilomycin A1 (Baf A1) (Hanhong, cat #AB23289) to inhibit vacuolation and maturation of notochordal cells, which can further affect the elongation of somite (*60*). The other group was treated with Cytochalasin D at a concentration of 1 μM. Another group was treated with DMSO as control group. All groups were treated for 4 h at 28°C and then embedded by 1% low melting point agarose for imaging.

#### Imaging of zebrafish embryos

The chorionic villi of target zebrafish embryos were peeled off with needle tips of 1 mL syringe. 1% low melting point agarose were melted at 65°C, for short-term imaging. The melted agarose was covered onto the embryos until cooling down to 26-30°C, and 0.3-0.5% low melting point agarose were used for long-term imaging to monitor the notochord maturation. During the agarose solidification, a blunted micropipette was used to place the embryo in a lateral orthotopic position. Then, the FRET data were collected by using Dragonfly Confocal Microscopy System (Seven-laser, 25× Silicon Oil (NA1.05)) (Dragonfly CR-DFLY-202-2540, U.S.A.). In addition, the embryos for phenotype were embedded into 3-5% methyl cellulose (Sigma-Aldrich), and the phenotypic images were taken by using Research Grade Stereomicroscope (Olympus SZX16, Japan).

---

**Fig. S1.**
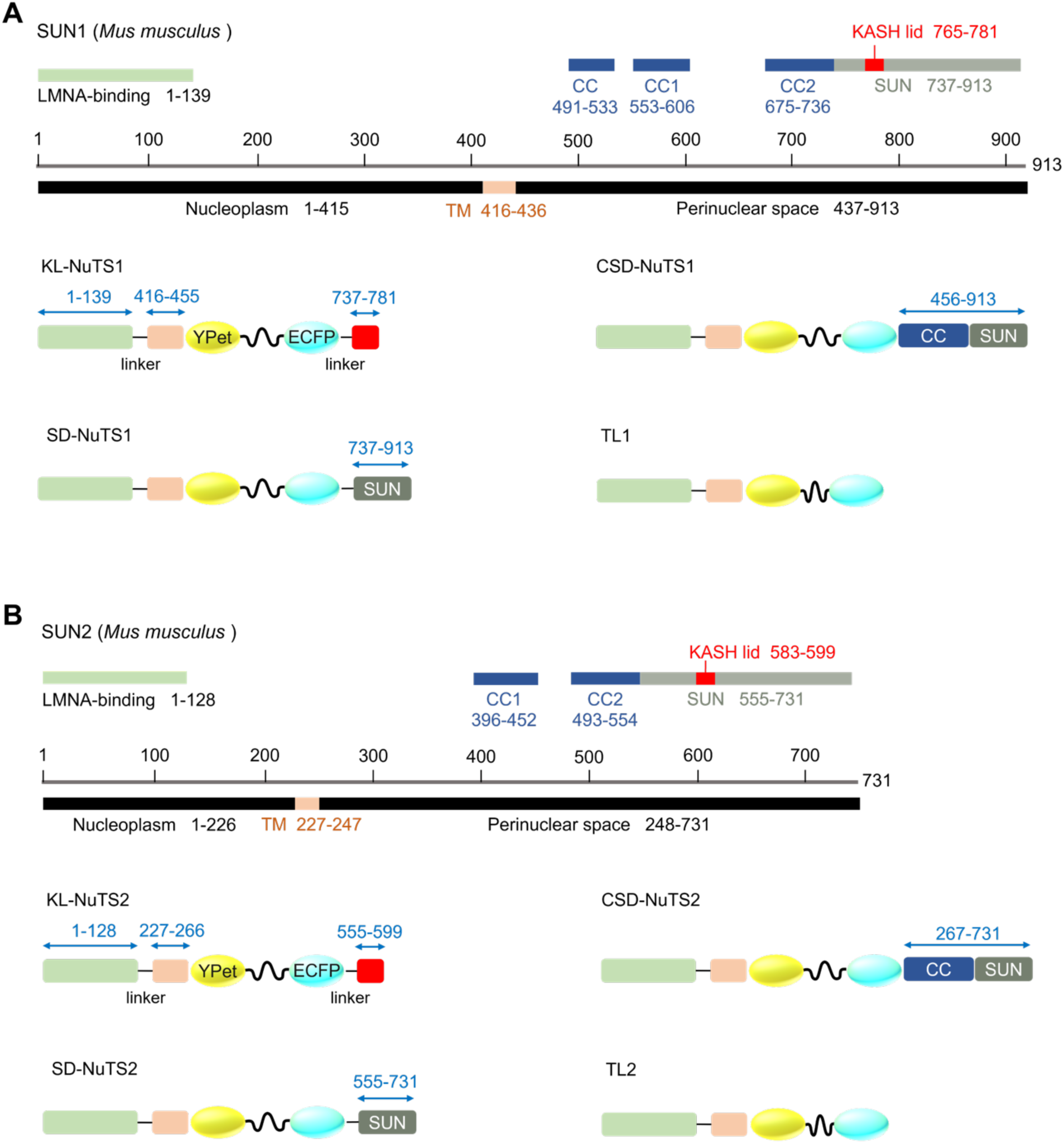
Designation of the FRET -based TL, KL-NuTS, SD-NuTS, and CSD-NuTS tension sensors. The sequence information was from National Library of Medicine, National Center for Biotechnology Information (Gene ID: 77053 for Sun1 Sad1 and UNC84 domain containing 2 [*Mus musculus* (house mouse)], and 223697 for Sun2 Sad1 and UNC84 domain containing 2 [*Mus musculus* (house mouse)]). The subdomain and amino acids sequence were from UniProt (Protein ID: Q9D666 for SUN domain-containing protein 1 and Q8BJS4 for SUN domain-containing protein 2). (**A**) for SUN1 and (**B**) for SUN2.

**Fig. S2.**
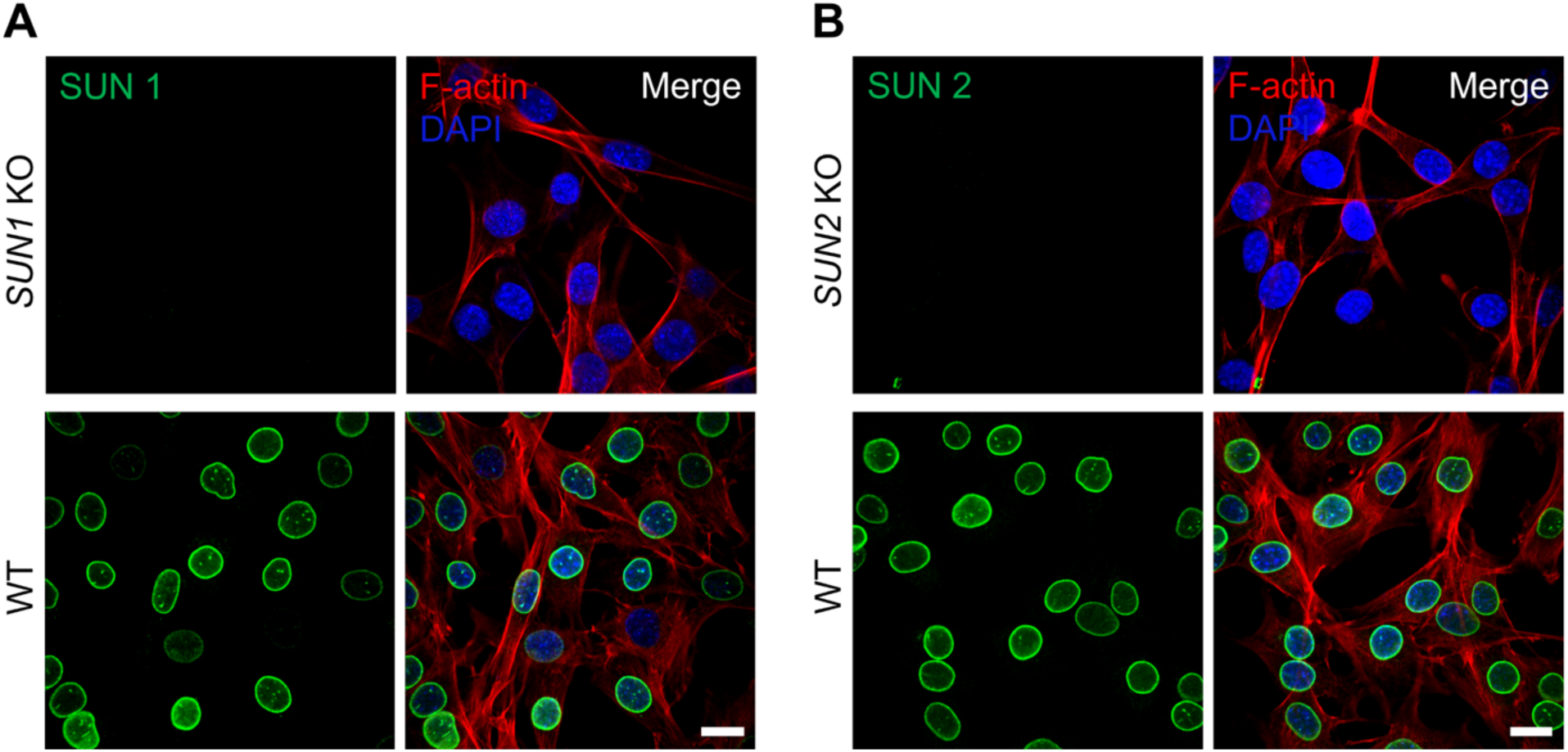
Identification of *SUN1* and *SUN2* knockout (KO) cell lines. (**A**) Representative immunostaining of SUN1 in *SUN1* KO and WT C2C12. Green for SUN1, red for F-actin, and blue for DAPI. Scale bars, 20 μm. (**B**) Representative immunostaining of SUN2 in *SUN2* KO and WT C2C12. Green for SUN2, red for F-actin, and blue for DAPI. Scale bars, 20 μm.

**Fig. S3.**
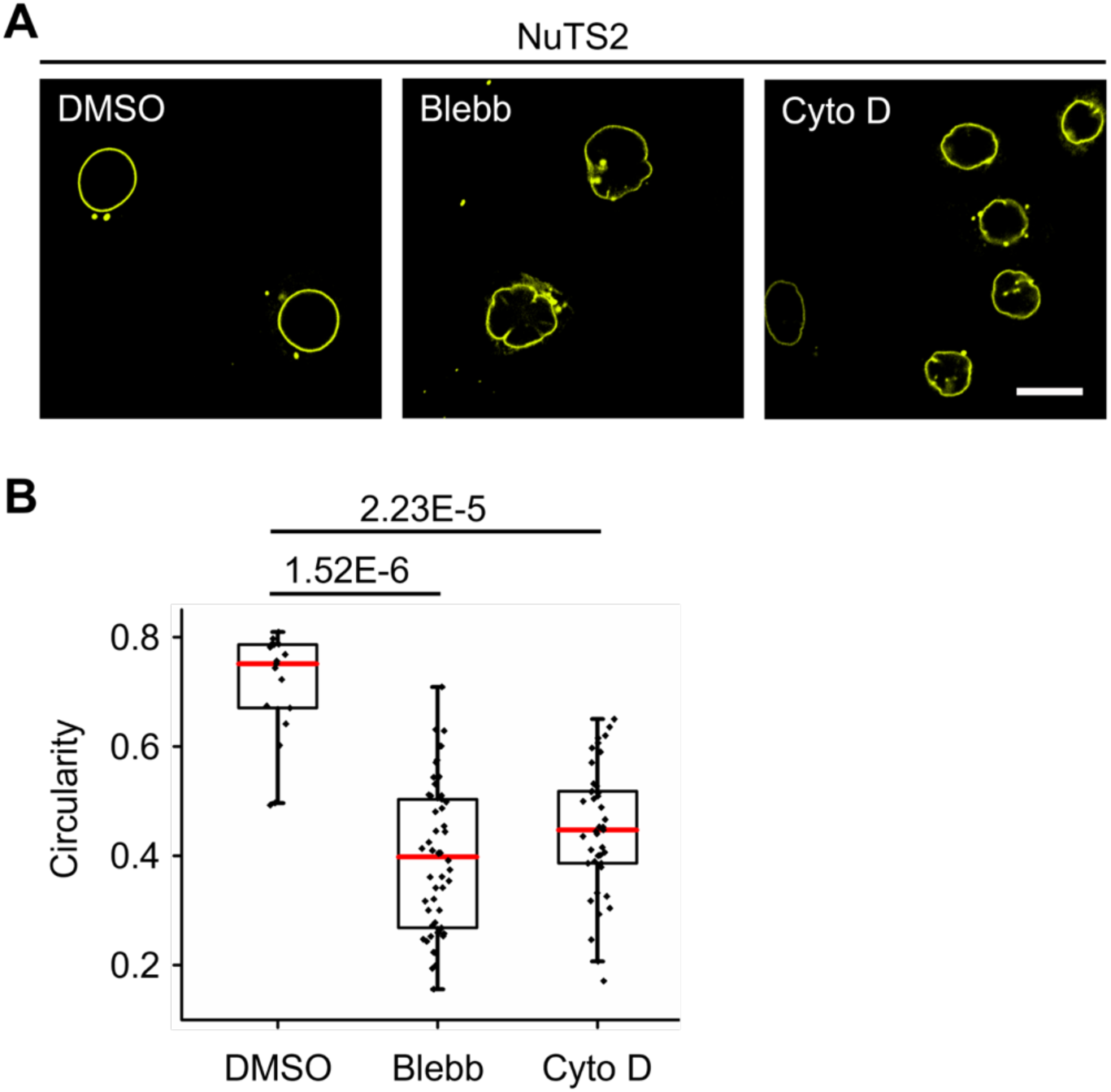
Nucleus morphology changes by drug treatment. (**A**) Representative nucleus shape images from HeLa cells with NuTS2 transiently expressed. Cells were treated with 1‰ DMSO for 30 min (control), 30 μM Blebbistatin (Blebb) for 1 h, and 1 μM Cytochalasin D (Cyto D) for 30 min, respectively. Yellow for YPet channel. Scale bars, 20 μm. (**B**) The quantified nuclear circularity (Mean ± SEM) of (**A**). The scatterplot represents the circularity in individual cells. n = 18, 53, 40. Three biological replicates. Ordinary one-way ANOVA Tukey’s multiple comparisons.

**Fig. S4.**
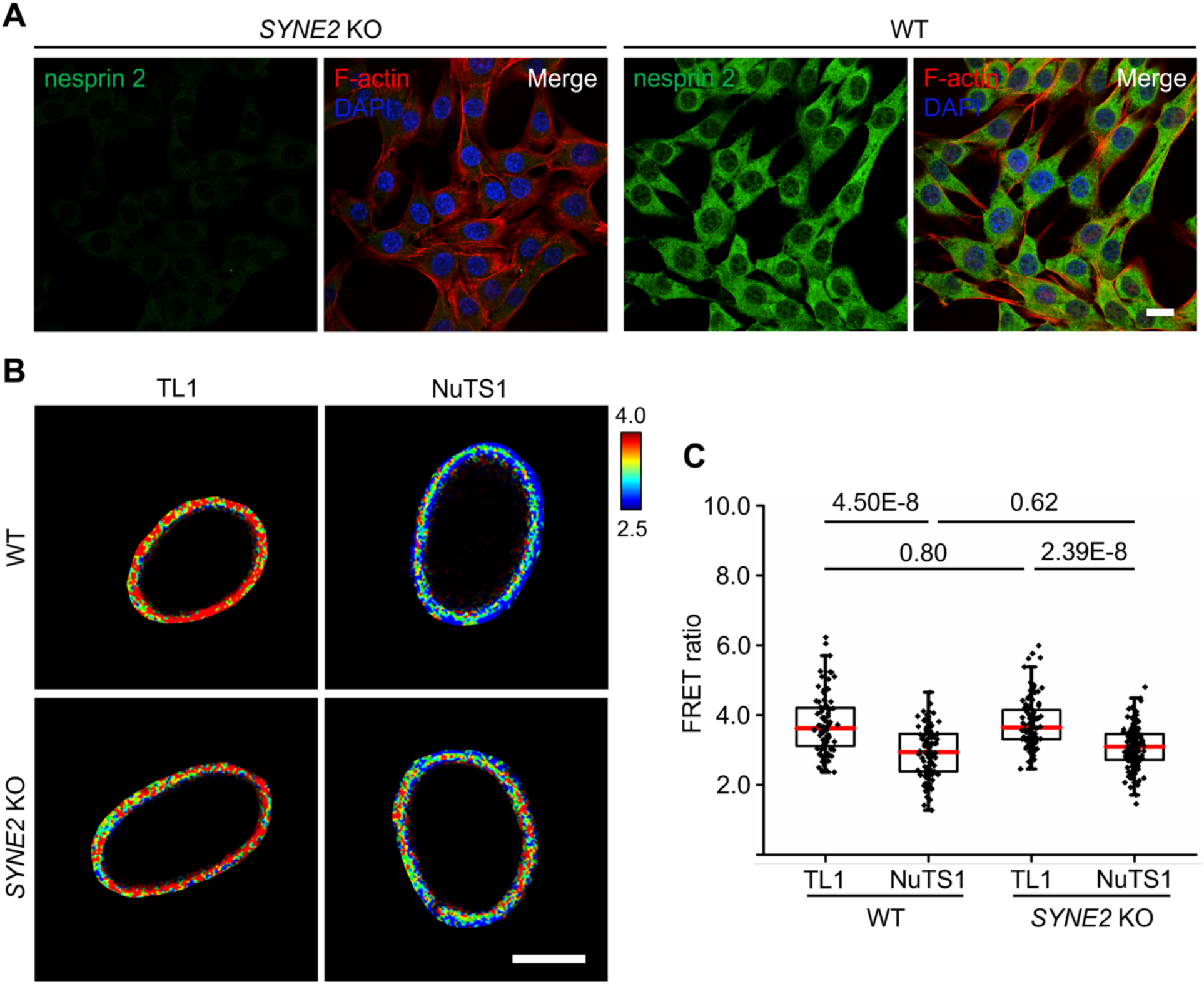
Identification of *SYNE2* knockout (KO) cell lines and responses of NuTS1 in *SYNE2* KO cells. (**A**) Representative immunostaining of nesprin 2 in *SYNE2* KO and WT C2C12. Green for nesprin 2, red for F-actin, and blue for DAPI. Scale bars, 20 μm. (**B**) Representative FRET ratiometric images of TL1 and NuTS1 expressed in *SYNE2* KO and WT C2C12, respectively. Scale bars, 10 μm. (**C**) The quantified FRET ratio (Mean ± SEM) of TL1 and NuTS1 from (**B**). The scatterplot represents the FRET/ECFP ratios in individual cells. n = 88, 108, 104, 133, three biological replicates. Ordinary one-way ANOVA Tukey’s multiple comparisons.

**Fig. S5.**
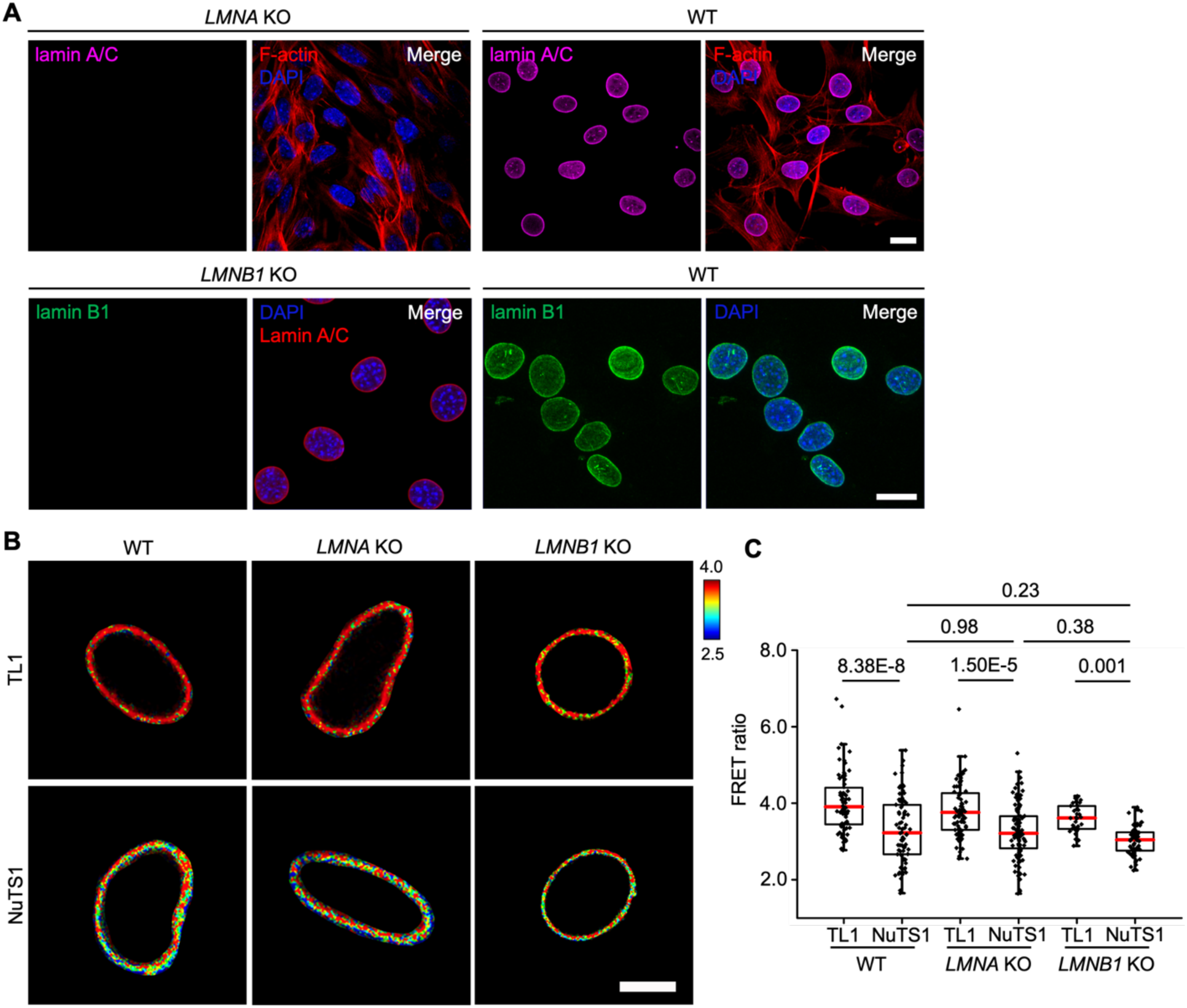
The responses of NuTS1 in *LMNA* knockout (KO) and *LMNB1* KO cells. (**A**) Representative immunostaining of lamin A/C in *LMNA* KO and WT C2C12. Magenta for lamin A/C, red for F-actin, and blue for DAPI. Representative immunostaining of lamin B1 in *LMNB1* KO and WT C2C12. Green for lamin B1, red for lamin A/C, and blue for DAPI. Scale bars, 20 μm. (**B**) Representative FRET ratiometric images of TL1 and NuTS1 expressed in *LMNA* KO, *LMNB1* KO and WT C2C12, respectively. Scale bars, 10 μm. (**C**) The quantified FRET ratio (Mean ± SEM) of TL1 and NuTS1 from (**B**). The scatterplot represents the FRET/ECFP ratios in individual cells. n = 71, 79, 84, 99, 38, 64, three biological replicates. Ordinary one-way ANOVA Tukey’s multiple comparisons.

**Fig. S6.**
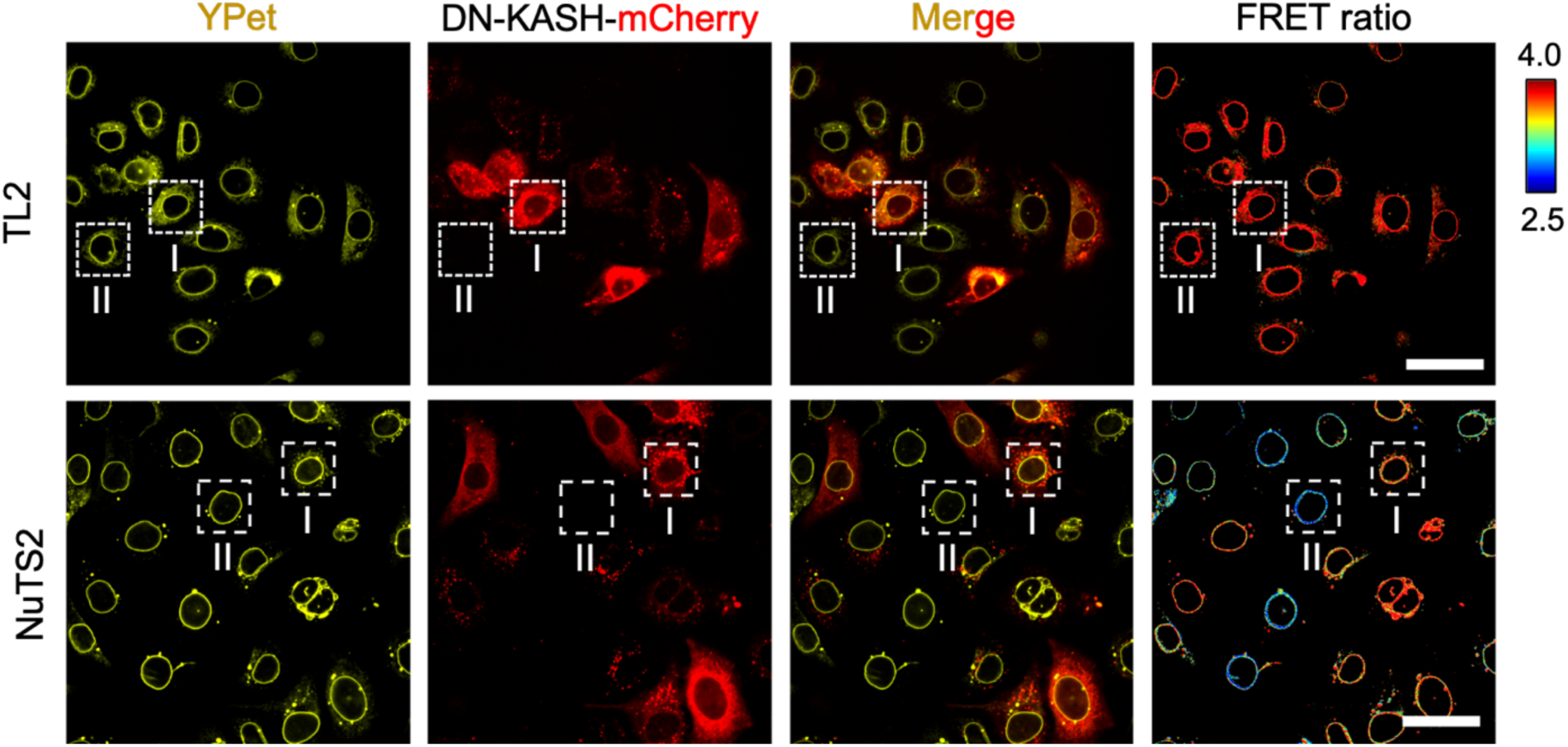
DN-KASH disrupts force transmission on NuTS2. Representative YPet intensity images, mCherry intensity images, and FRET ratiometric images from TL2 and NuTS2 stable HeLa cells with DN-KASH transiently expressed for 36 h, respectively. Yellow for YPet and red for DN-KASH-mCherry. The region I represents the cell with DN-KASH expressing, and the region II represents the cell without DN-KASH. Scale bars, 50 μm.

**Fig. S7.**
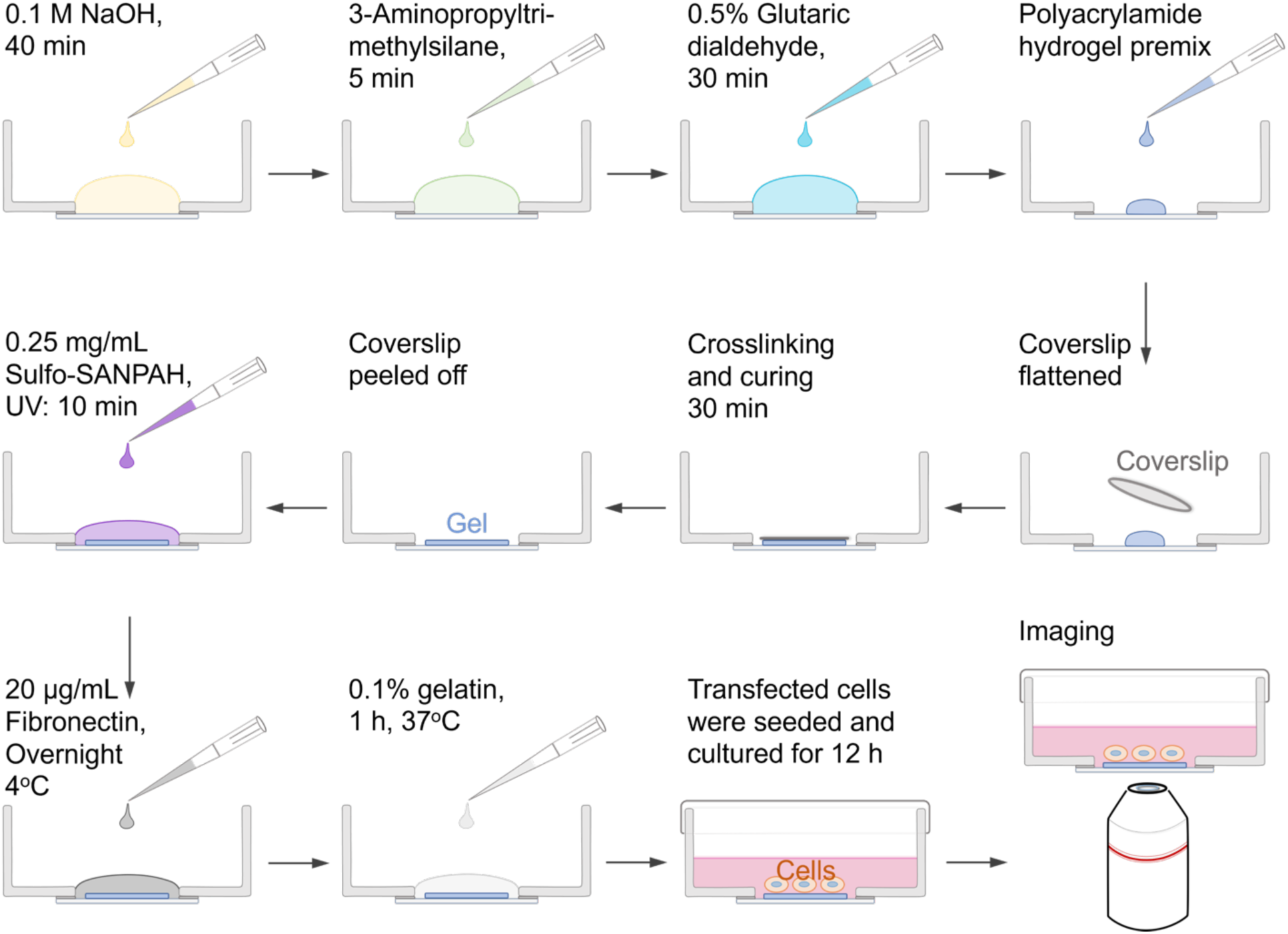
The schematic of PAA hydrogel preparation with different stiffness. The processes of gel preparation and cell seeding. Glass bottom dishes were treated with following NaOH, 3-Aminopropyltri-methylsilane, Glutaric dialdehyde to make the surface ready for gel adhesion. Then, the polyacrylamide hydrogel premix was dropped and covered with a coverslip rapidly, and the coverslip was peeled off after gel cured. Next, the gel was immersed into the crosslinker (Sulfo-SANPAH) and coated with Fibronectin and gelatin. Finally, transfected cells were seeded onto the gels for cell culture and imaging.

**Fig. S8.**
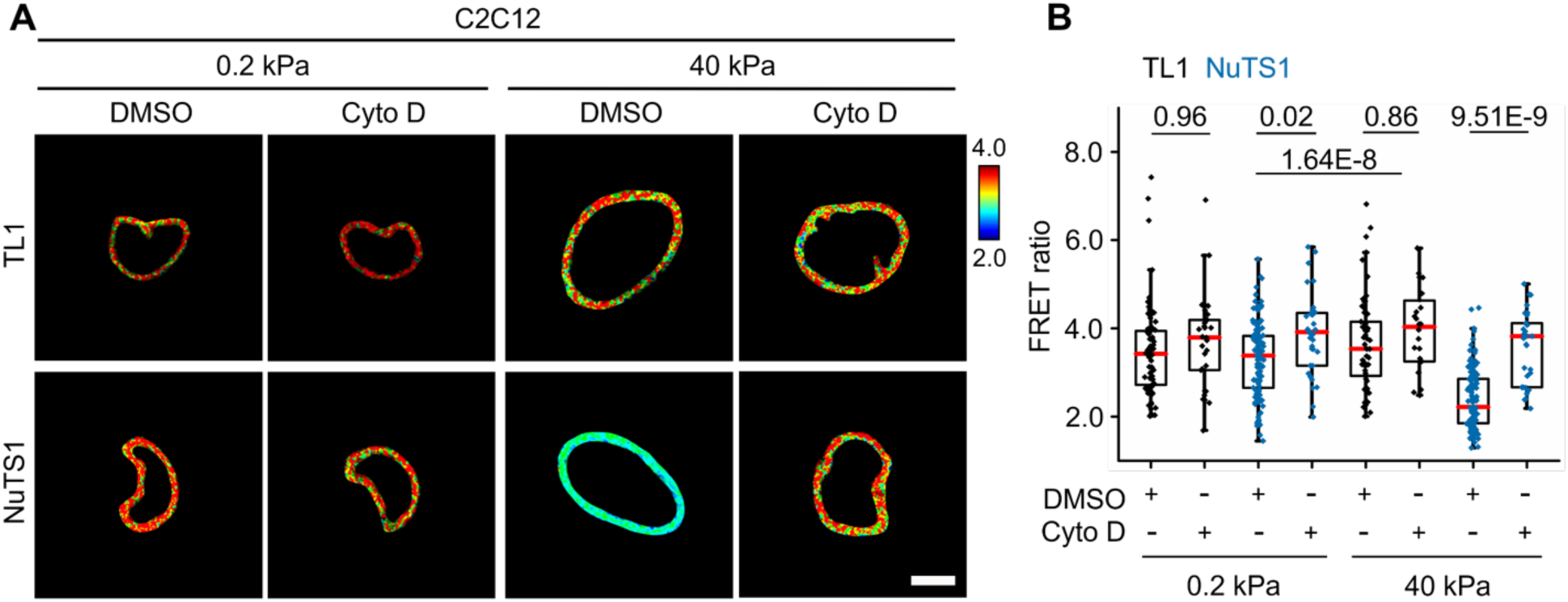
The responses of NuTS1 to the matrix stiffness. (**A**) Representative FRET ratiometric images of TL1 and NuTS1 in WT C2C12 cells that seeded on fibronectin coated soft (0.2 kPa) and stiff (40 kPa) PAA gel, respectively. Scale bars, 10 μm. (**B**) The quantified FRET ratio (Mean ± SEM) of TL1 and NuTS1 from (**A**). The scatterplot represents the FRET/ECFP ratios in individual cells. n = 79, 32, 129, 39, 64, 27, 130, 34, two biological replicates, respectively. Ordinary one-way ANOVA Tukey’s multiple comparisons.

**Fig. S9.**
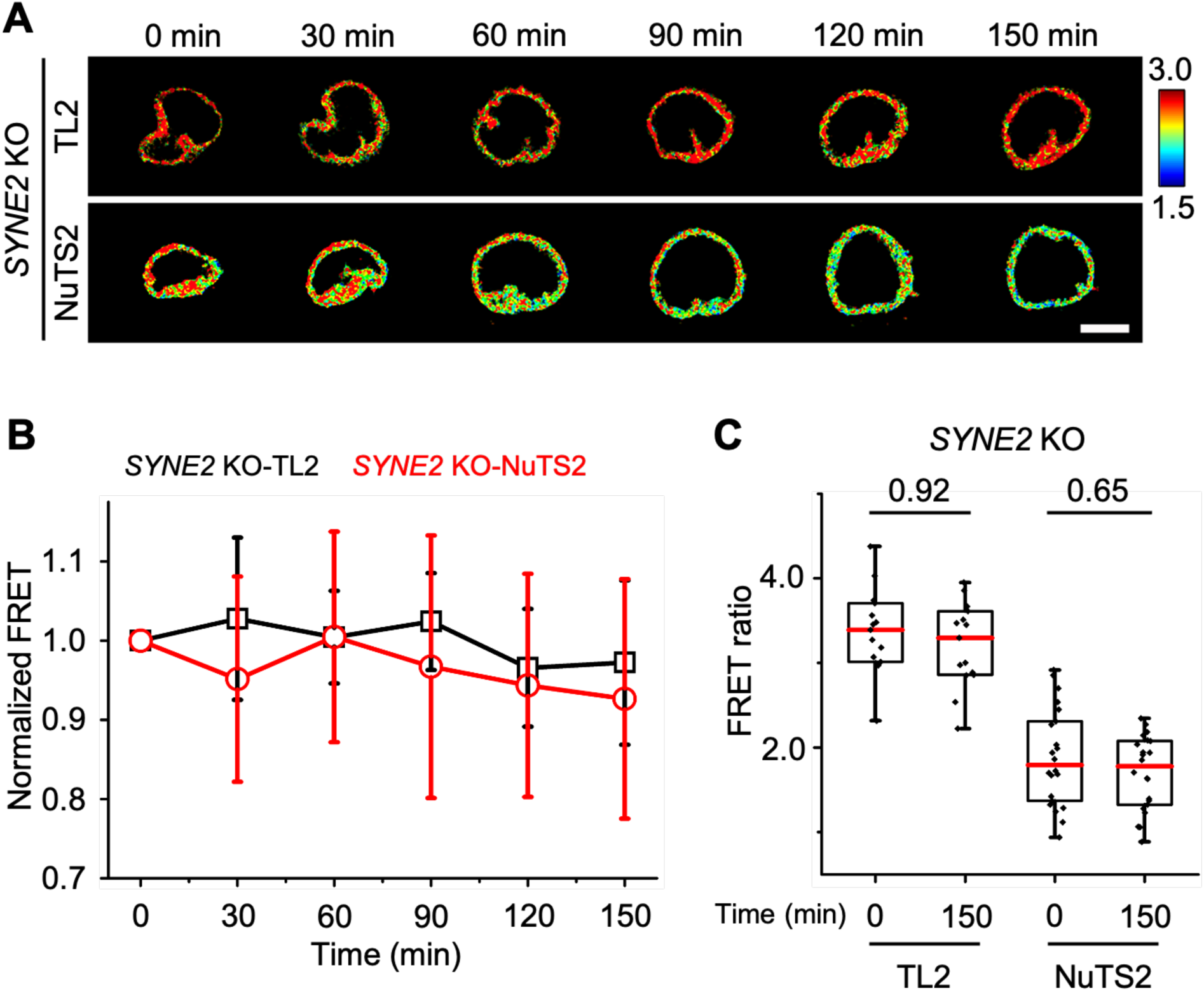
FRET dynamics of NuTS2 during cell adhesion in *SYNE2* KO cell. (**A**) Representative FRET ratiometric images of TL2 and NuTS2 expressed in *SYNE2* KO C2C12 cells that seeded on glass bottom dish. The images were taken with 30 min interval during cell adhesion for 2.5 h. Scale bars, 10 μm. (**B**) The normalized FRET ratio (Mean ± SEM) of TL2 and NuTS2 from (**A**). n = 15, 22, three biological replicates. (**C**) The quantified FRET ratio (Mean ± SEM) of TL2 and NuTS2 before (0 min) and after (150 min). The scatterplot represents the FRET/ECFP ratios in individual cells. n = 15, 15, 22, 22, three biological replicates. Ordinary one-way ANOVA Tukey’s multiple comparisons.

**Fig. S10.**
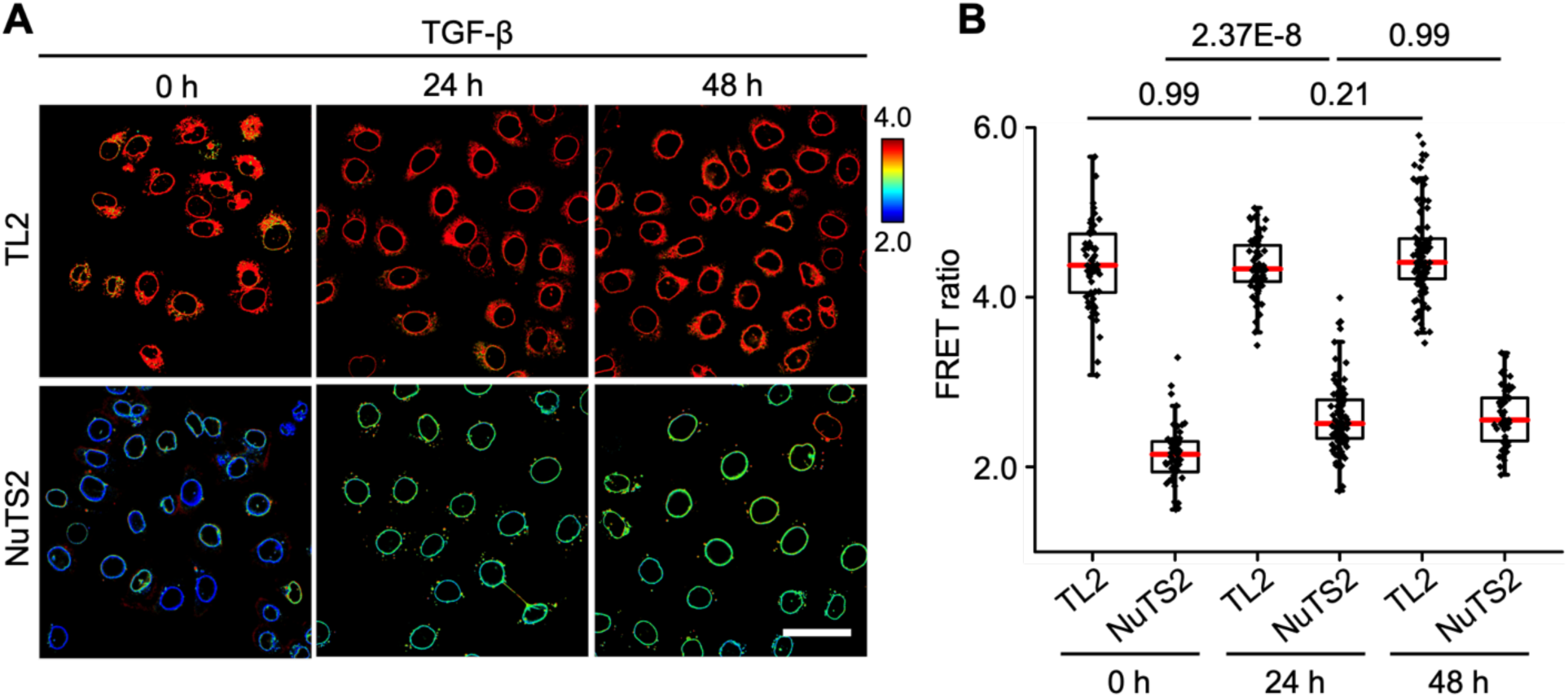
The responses of NuTS1 during EMT in HeLa cells. (**A**) Representative image of TL2 and NuTS2 stable HeLa cells subjected to EMT induction by treating with 5 ng/mL TGF-beta1 growth factor for 24 h and 48 h, respectively. Scale bars, 50 μm. (**B**) The quantified FRET ratio (Mean ± SEM) of TL2 and NuTS2 from (**A**). The scatterplot represents the FRET/ECFP ratios in individual cells. n = 64, 64, 79, 120, 124, 64. Three biological replicates. Ordinary one-way ANOVA Tukey’s multiple comparisons.

**Fig. S11.**
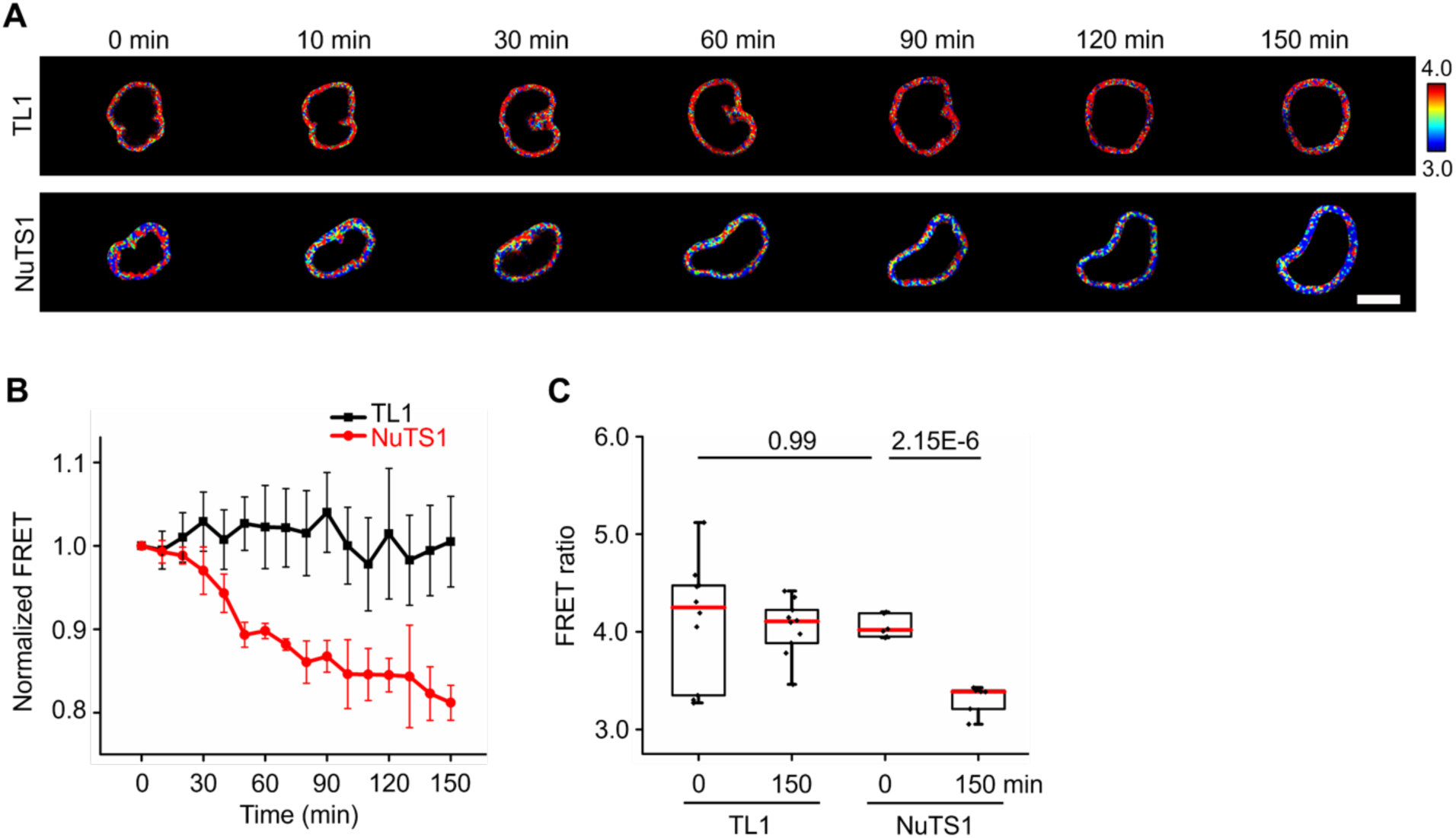
The mechanical forces transmit across SUN1 during cell spreading by NuTS1. (**A**) Representative FRET ratiometric images of TL1 and NuTS1 expressed in WT C2C12 cells on glass bottom dish. The images were taken with 10 min interval during the first 3 h of cell adhesion. Scale bars, 10 μm. (**B**) The normalized FRET ratio (Mean ± SEM) of TL1 and NuTS1 from (**A**). n = 10, 6. Three biological replicates. (**C**) The quantified FRET ratio (Mean ± SEM) of TL1 and NuTS1 before (0 min) and after (150 min), the scatterplot represents the FRET/ECFP ratios in individual cells. n = 10, 10, 6, 6. Three biological replicates. Ordinary one-way ANOVA Tukey’s multiple comparisons.

**Fig. S12.**
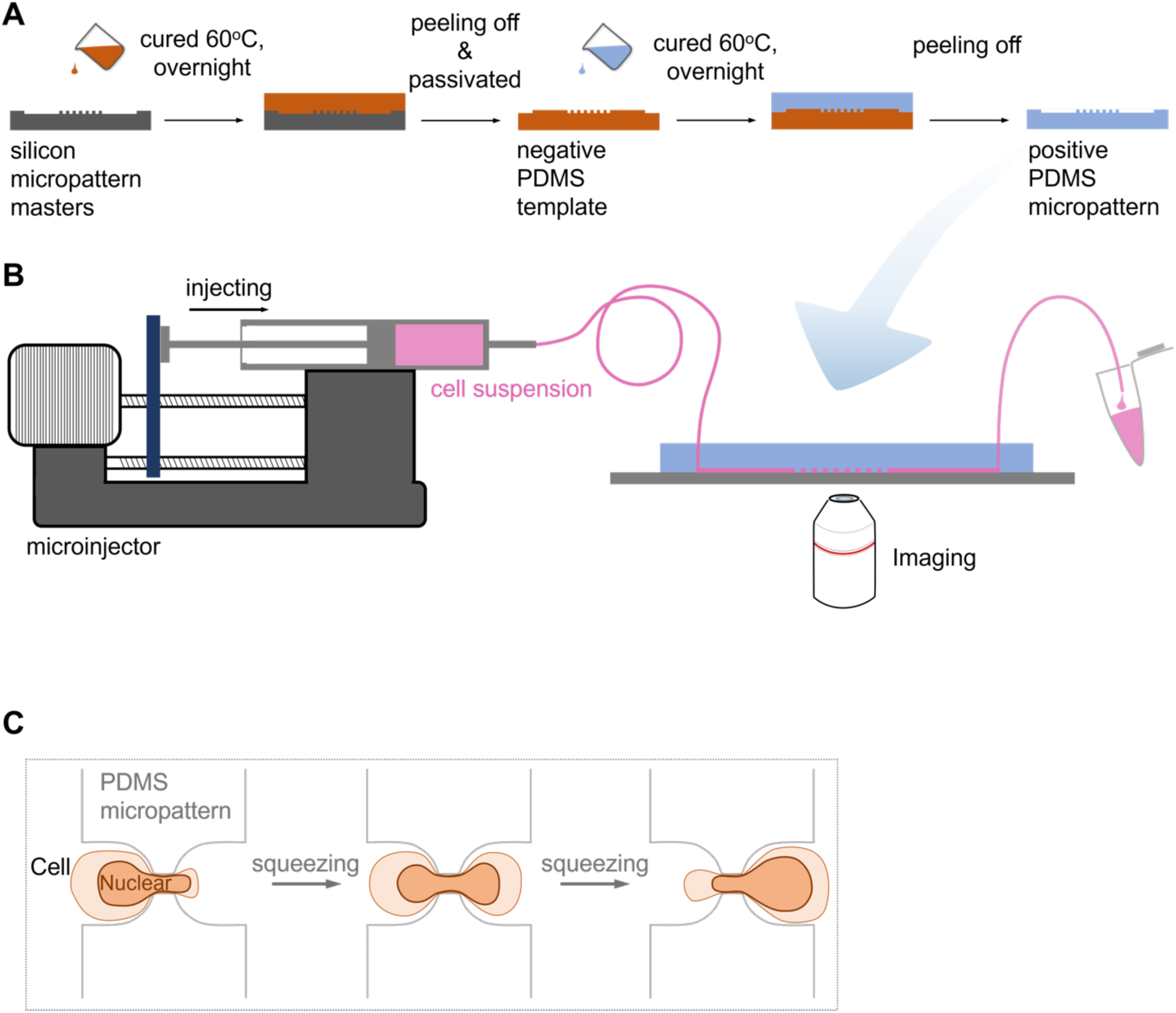
The schematics of squeezing model preparation and its working mechanism. (**A**) The schematics of squeezing model preparation. The micropattern was prepared with 10: 1 PDMS by using twice-guided molds from a fabricated silicon master. (**B**) The processes of cell loading by using the microinjector. The Hela stable cells were detached, filtrated by flow cytometric sieve and loaded into the squeezing model, and the flow rate was 0.002 mL/min. (**C**) Schematic diagram of nuclear deformation during squeezing.

**Fig. S13.**
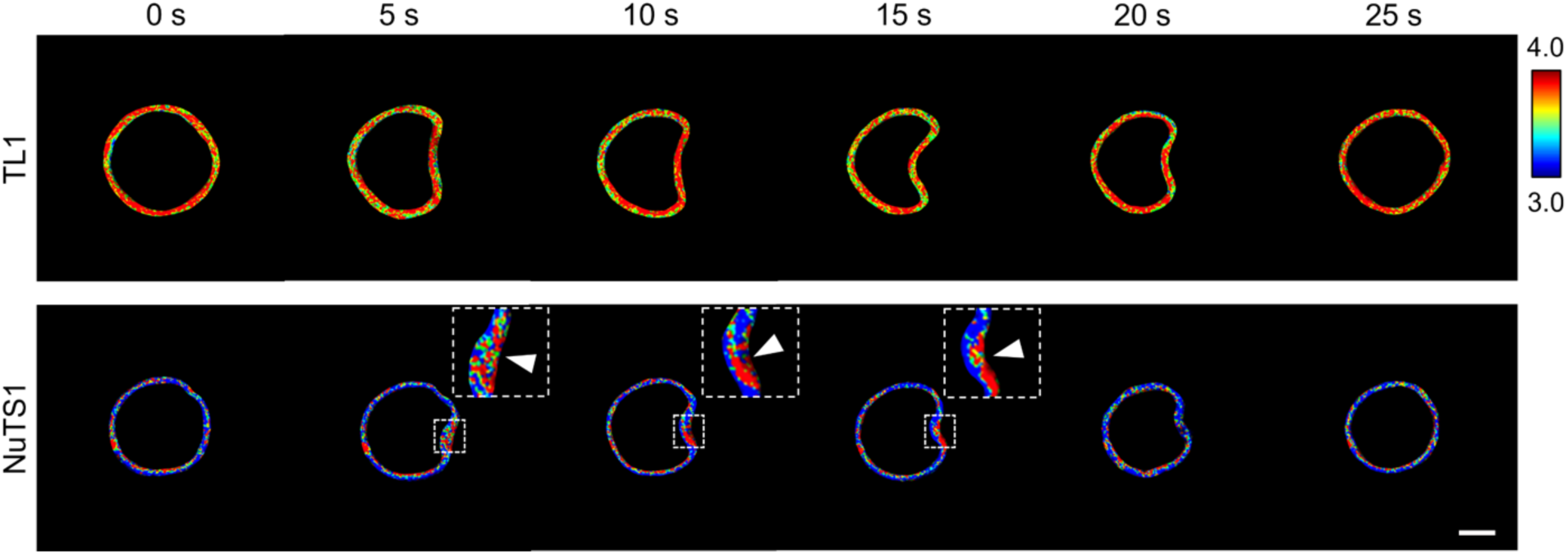
The dynamic changes of FRET ratio with local force stimuli at nuclear membrane. The dynamic FRET ratio changes with the continuous pressing of micropipette at local region. Representative FRET ratiometric images in Hela with TL1 and NuTS1 transfected. White arrows show the local force stimuli. Scale bars, 10 μm.

**Fig. S14.**
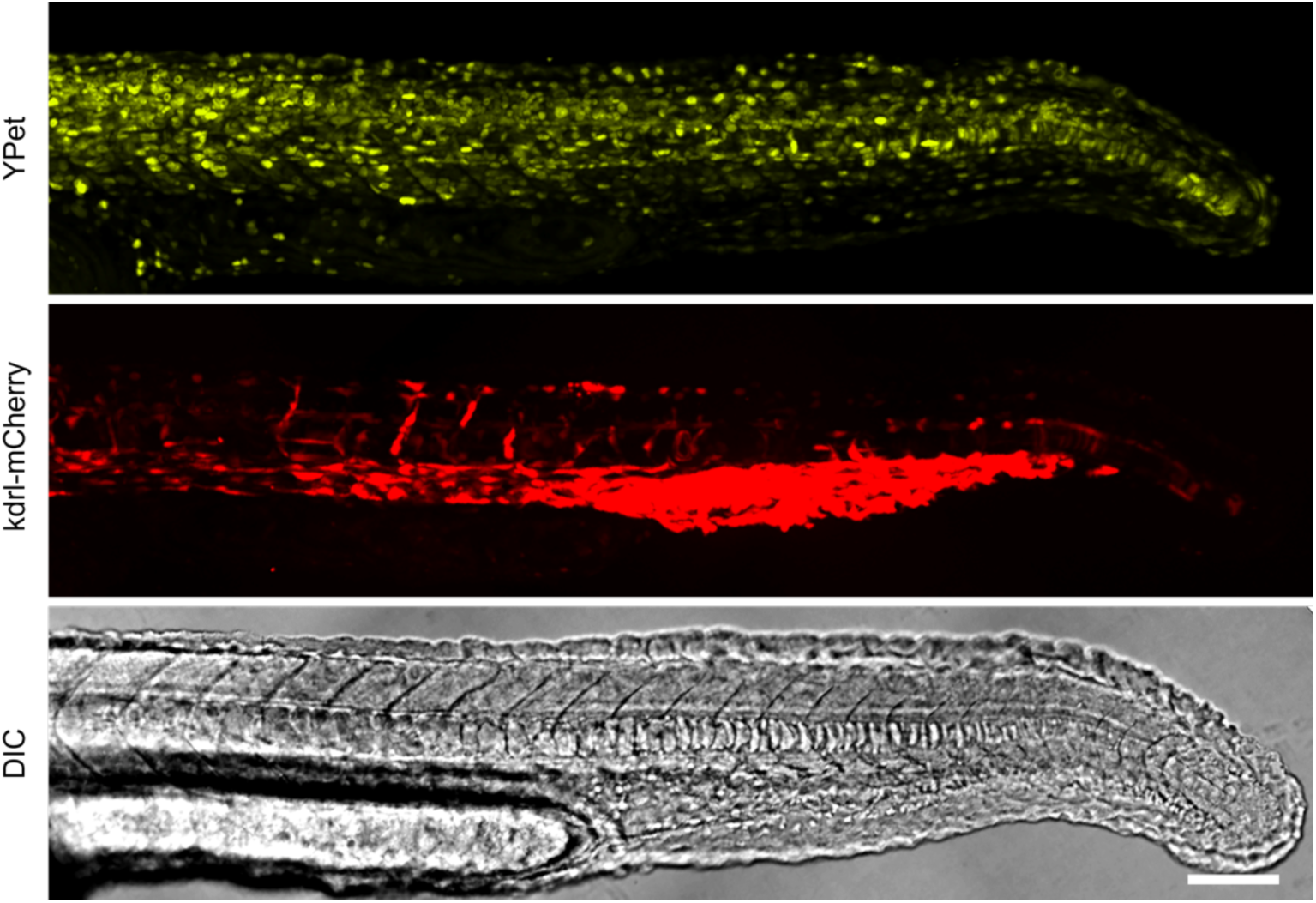
The expression of *Kdrl:mCherry* in zebrafish. Representative images of NuTS2 and *kdrl*:mCherry expressed in zebrafish at 24 hpf. Scale bars, 100 μm.

**Fig. S15.**
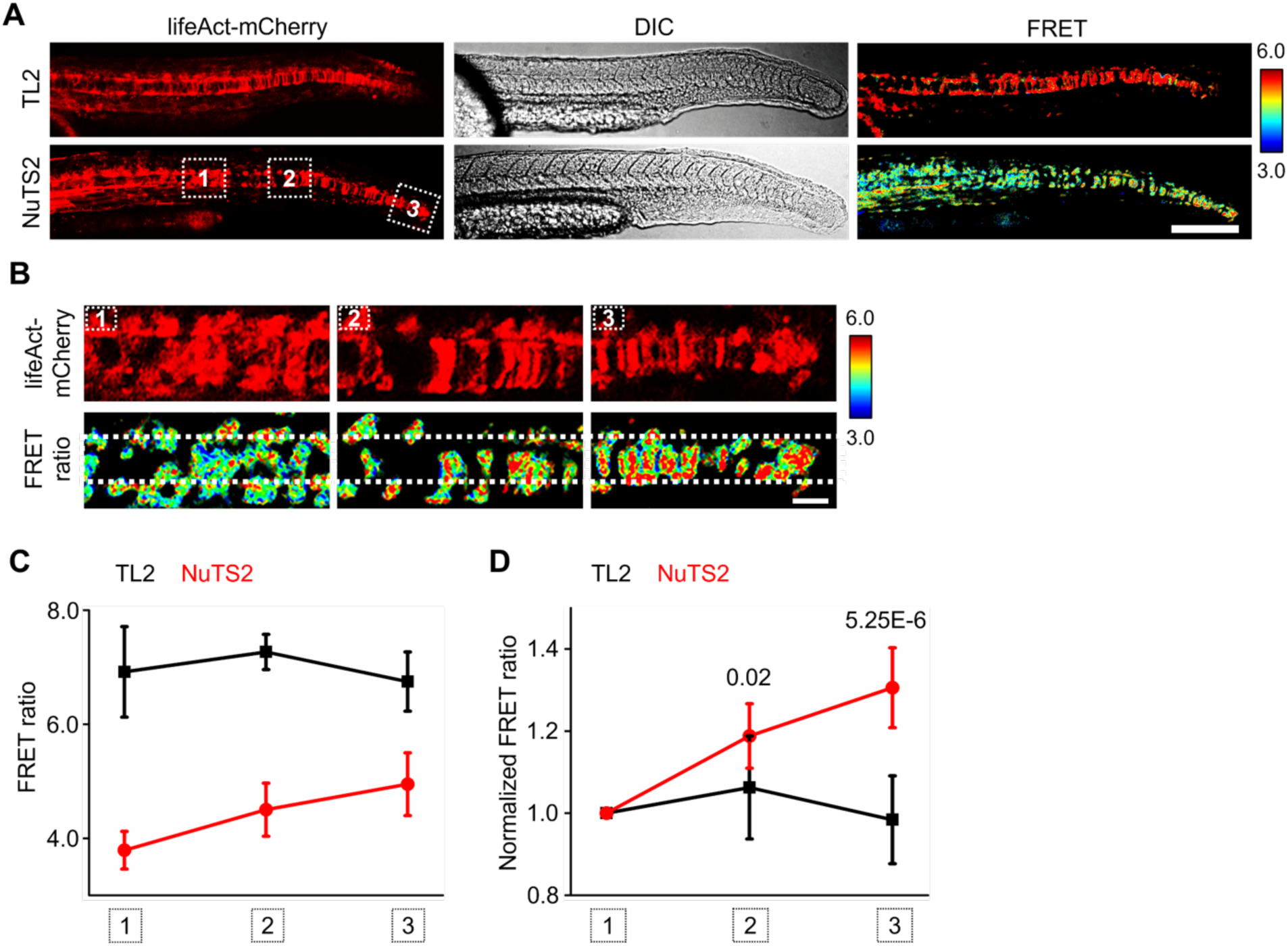
**The tension forces measured by NuTS2 along notochord in zebrafish embryos**. (**A**) Representative FRET ratiometric images of TL2 or NuTS2 and lifeAct-mCherry co-expressed in zebrafish, respectively. The TL2 or NuTS2 mRNA and lifeAct-mCherry mRNA were injected into embryo at 1-2 cell stage, then the expression of mRNA at the notochord in fishtail was tested by confocal microscopy. Scale bars, 200 μm. (**B**) The amplification represents the zoom in images of the marked location from (**A**). Scale bars, 20 μm. (**C**) The quantified FRET ratio (Mean ± SEM) of TL2 and NuTS2 at different locations from (**B**). The average of FRET ratio from three regions was calculated for each localization. n = 8, 10 for TL2 and NuTS2, three biological replicates, respectively. Ordinary one-way ANOVA Tukey’s multiple comparisons. (**D**) The normalized FRET ratio (Mean ± SEM) of TL2 and NuTS2 at different locations from (**C**). n = 8, 10 for TL2 and NuTS2, three biological replicates, respectively. Ordinary one-way ANOVA Tukey’s multiple comparisons.

**Fig. S16.**
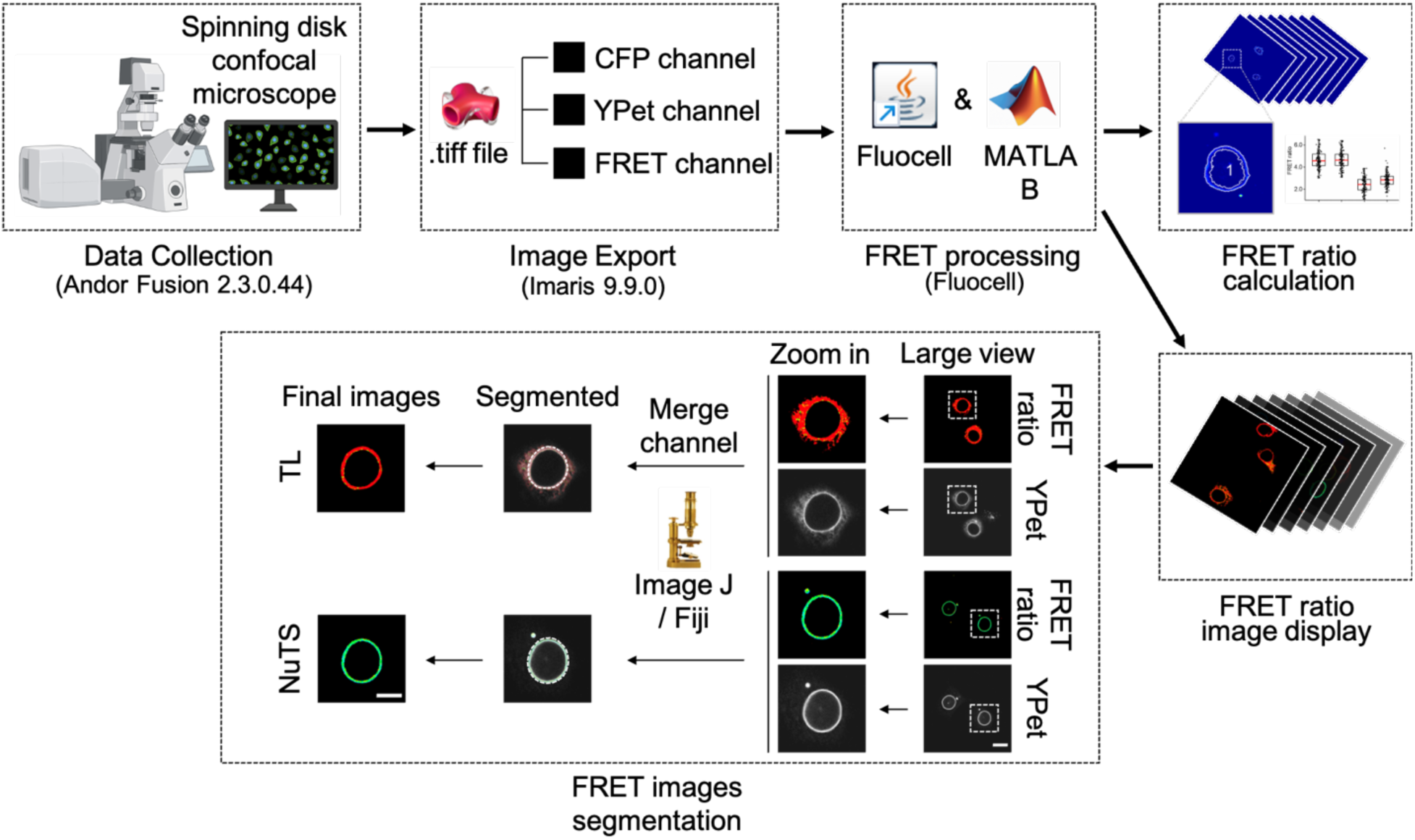
Procedures for FRET imaging acquisition and data analysis. The data was collected by confocal microscope and exported by using Imaris software, then both the FRET ratio calculation and FRET ratio images were processed by Fluocell software. The final FRET ratio images were obtained through segmentation by Image J/Fiji.

**Table S1.**
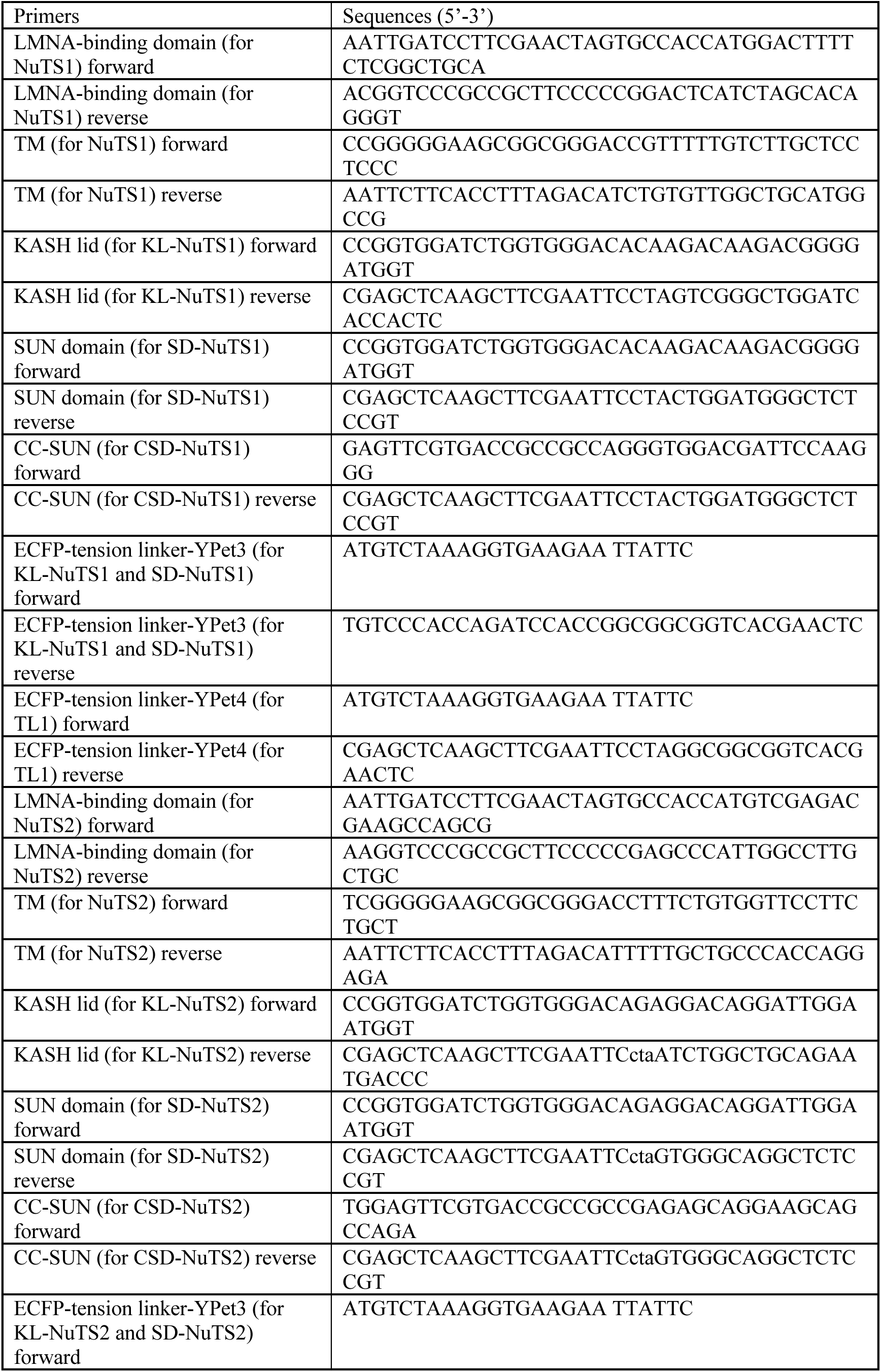
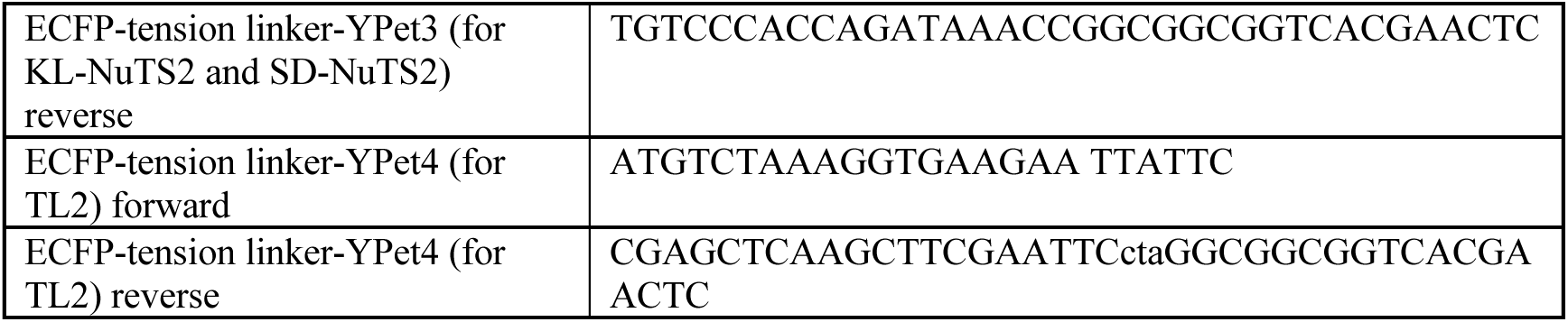
The primers for tension sensor construction.

**Movie S1.**

The dynamic FRET changes of TL2 and NuTS2 in HeLa stable cells with Cyto D treatment.

**Movie S2.**

The dynamic FRET changes of TL2 and NuTS2 in HeLa stable cells with Y-27632 treatment.

